# Fetally-encoded GDF15 and maternal GDF15 sensitivity are major determinants of nausea and vomiting in human pregnancy

**DOI:** 10.1101/2023.06.02.542661

**Authors:** M Fejzo, N Rocha, I Cimino, SM Lockhart, C Petry, RG Kay, K Burling, P Barker, AL George, N Yasara, A Premawardhena, S Gong, E Cook, K Rainbow, DJ Withers, V Cortessis, PM Mullin, KW MacGibbon, E Jin, A Kam, A Campbell, O Polasek, G Tzoneva, FM Gribble, GSH Yeo, BYH Lam, V Saudek, IA Hughes, KK Ong, JRB Perry, A Sutton Cole, M Baumgarten, P Welsh, N Sattar, GCS Smith, DS Charnock Jones, AP Coll, CL Meek, S Mettananda, C Hayward, N Mancuso, S O’Rahilly

## Abstract

Human pregnancy is frequently accompanied by nausea and vomiting that may become severe and life-threatening, as in hyperemesis gravidarum (HG), the cause of which is unknown. Growth Differentiation Factor-15 (GDF15), a hormone known to act on the hindbrain to cause emesis, is highly expressed in the placenta and its levels in maternal blood rise rapidly in pregnancy. Variants in the maternal *GDF15* gene are associated with HG. Here we report that fetal production of GDF15, and maternal sensitivity to it, both contribute substantially to the risk of HG. We found that the great majority of GDF15 in maternal circulation is derived from the feto-placental unit and that higher GDF15 levels in maternal blood are associated with vomiting and are further elevated in patients with HG. Conversely, we found that lower levels of GDF15 in the non-pregnant state predispose women to HG. A rare C211G variant in *GDF15* which strongly predisposes mothers to HG, particularly when the fetus is wild-type, was found to markedly impair cellular secretion of GDF15 and associate with low circulating levels of GDF15 in the non-pregnant state. Consistent with this, two common *GDF15* haplotypes which predispose to HG were associated with lower circulating levels outside pregnancy. The administration of a long-acting form of GDF15 to wild-type mice markedly reduced subsequent responses to an acute dose, establishing that desensitisation is a feature of this system. GDF15 levels are known to be highly and chronically elevated in patients with beta thalassemia. In women with this disorder, reports of symptoms of nausea or vomiting in pregnancy were strikingly diminished. Our findings support a causal role for fetal derived GDF15 in the nausea and vomiting of human pregnancy, with maternal sensitivity, at least partly determined by pre-pregnancy exposure to GDF15, being a major influence on its severity. They also suggest mechanism-based approaches to the treatment and prevention of HG.

## Main

Nausea and vomiting affects approximately 70% of human pregnancies and can often be debilitating [1]. Hyperemesis gravidarum (HG) is diagnosed when nausea and vomiting are so severe that women are unable to eat and/or drink normally and have greatly limited daily activity. This is frequently accompanied by weight loss and electrolyte disturbance which can carry significant risks to the longer-term health of both mother and offspring [1]. In the USA, HG is the leading cause of hospitalization in early pregnancy and the 2^nd^ most common cause of pregnancy hospitalization overall [2]. Until recently there has been no significant advance in the understanding of the molecular pathogenesis of nausea and vomiting of pregnancy (NVP) or HG. A body of evidence implicating GDF15, a circulating member of the TGF-ß superfamily, in these disorders has been emerging. In the non-pregnant state, GDF15 is ubiquitously produced in response to a range of cellular stresses. Its receptor, a heterodimer of GFRAL and RET, is only expressed in the hindbrain where its activation leads to nausea, vomiting, and aversive responses. For example, cis-platinum therapy acutely elevates circulating GDF15 and the vomiting that occurs as a result of this is, in non-human primates, largely prevented by neutralising GDF15 [3]. The presence of high levels of GDF15 (then called MIC-1) in maternal blood in normal human pregnancy was first reported in 2000 [4] by Breit and colleagues who first described the hormone. Recently, GDF15 was found to be one of the most abundant peptides secreted from human trophoblast organoids [5] and *GDF15* mRNA is more abundant in placental mRNA than in all the tissues examined by the GTEx consortium [6]. When compared to women who had low levels of nausea or vomiting, concentrations of GDF15 in maternal circulation have been reported to be higher in women experiencing vomiting in pregnancy [7] and in a small group of women with HG [8]. The notion that GDF15 may have a primary role in the aetiology of HG, rather than increase as a consequence of the condition, was supported by the findings of the first genome wide association study of women with HG, which reported several independent variants close to the *GDF15* gene as the most highly associated SNPs in the maternal genome [9]. Subsequently, Fejzo et al undertook an exome sequencing study in HG cases and controls and found that a rare, heterozygous missense variant in GDF15 (C211G) was highly enriched in HG cases vs controls [10]. However, to date, a mechanistic basis for these genetic associations has not been clearly elucidated. Here we demonstrate that GDF15 is truly elevated in NVP and HG and that the vast majority of GDF15 is of fetal origin. Remarkably, we show that rare and common genetic variants in *GDF15* that increase HG risk do so by lowering circulating pre-pregnancy GDF15, and that women with conditions that increase GDF15 in the non-pregnant state are protected from NVP/HG; findings which appear to conflict with the known anorectic and emetic actions of GDF15. We resolve this apparent paradox by demonstrating that the anorectic actions of the GDF15-GFRAL axis are subject to desensitization and propose that antecedent levels of GDF15 influence maternal sensitivity to the surge of fetal derived GDF15 which occurs from early pregnancy, thus determining the pregnant woman’s susceptibility to develop NVP and HG.

### Circulating GDF15 levels are elevated in mothers with either Hyperemesis Gravidarum or vomiting in pregnancy

A common genetic variant at amino acid 202 of GDF15 (H to D, hereafter H202D) has recently been shown to systematically and substantially interfere with measurements of the peptide by reagents used in most of the studies that have reported GDF15 concentrations in human circulation [11]. We therefore commenced our investigations by measuring GDF15 in blood using an immunoassay that is less susceptible to confounding by the H202D variant (**Supplementary Table 1**); samples were taken at ∼15 weeks gestation from women who completed a questionnaire relating to NVP. GDF15 levels were significantly increased in women reporting vomiting compared to those reporting no nausea or vomiting (**Figure 1A and Supplementary Table 2-3**). In a second study, we obtained blood samples from 57 women presenting to hospital with HG and from 56 controls who reported low levels of nausea or vomiting. Participants in each group were of similar age and BMI and were predominantly in first trimester of pregnancy when recruited (**Supplementary Table 4**). GDF15 levels (measured by an assay that is not susceptible to interference by H202D [11]) were significantly higher in women with HG vs those without (**Figure 1B and C, Supplementary Table 5**). These results increase confidence that there is a true association between maternal GDF15 levels with HG and levels of vomiting in pregnancy.

**Figure 1.**
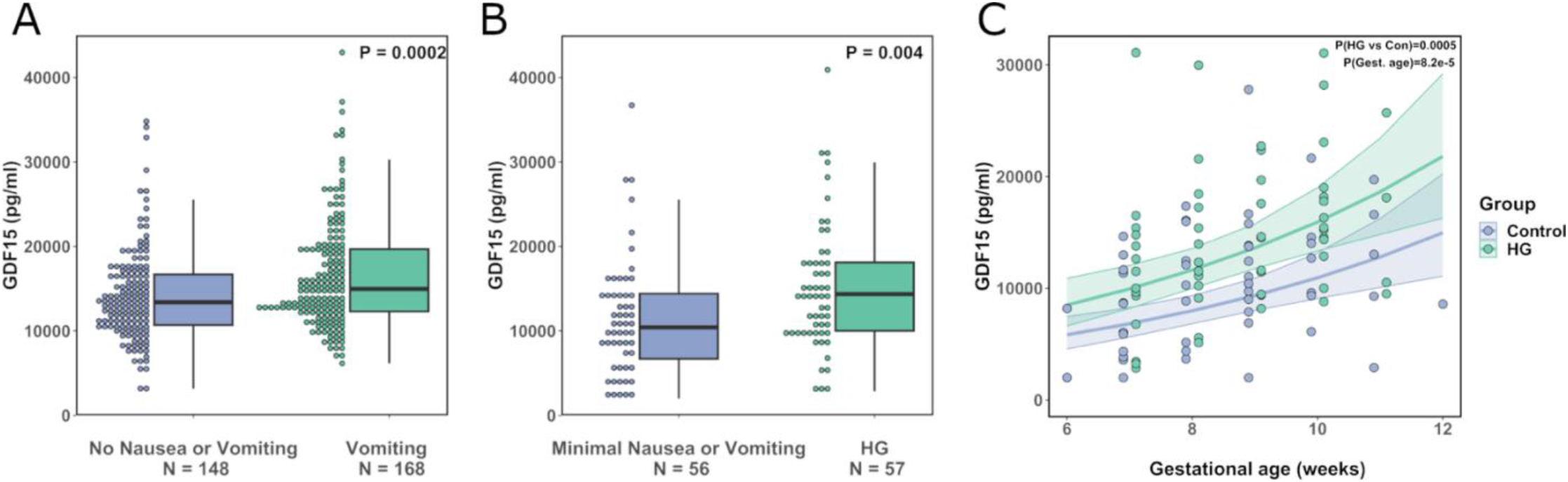
Circulating GDF15 is elevated in women experiencing nausea and vomiting in pregnancy and hyperemesis gravidarum. **A:** Dot and box plots illustrating the distribution of circulating GDF15 levels in women of ∼15 weeks’ gestation with a history of vomiting in pregnancy vs those reporting no nausea and vomiting in pregnancy. P-value is from an unadjusted linear regression model using natural-log transformed GDF15 concentrations. **B:** Dot and box plots illustrating the distribution of GDF15 levels (Mean gestational age ∼ 10 weeks) in women presenting with hyperemesis gravidarum (HG) and those with low levels of nausea and vomiting in pregnancy. P-value is from an unadjusted linear regression model using natural-log transformed GDF15 concentrations. **C:** Scatter plot illustrating the relationship between gestational age and GDF15 in the first trimester. The trend lines show predicted values of GDF15 levels (+/-95% CI) in women with and without HG from a linear regression model of natural-log transformed GDF15 with gestational age and HG status included as predictor variables. The P-values are derived from the same regression model for the effect of HG (HG vs Con) and gestational age (Gest. Age). 5 participants (HG = 1, Control = 4) included in the analysis in panel (B) are not plotted or included in this model as they were recruited after the first trimester.

### GDF15 in the maternal circulation is of predominantly fetal origin

GDF15 is widely expressed and, although the placenta is a site of high levels of expression, the relative contribution of the fetal and maternal tissues has not been established. To examine this, we developed mass spectrometry-based assays capable of distinguishing between GDF15 carrying a histidine or an aspartate at position 202 (position 6 in the mature circulating molecule) (**Extended Data Figure 1, Figure 2A**). Using placental RNAseq data and maternal DNA from the Pregnancy Outcome Prediction (POP) study cohort [6] we genotyped offspring and mothers (**Supplementary Table 6**) and studied 7 H202D discordant offspring/mother pairs in which either the fetus or the mother alone was heterozygous at this site. Strikingly, in maternal plasma where the mother was heterozygous at H202D the discordant maternal peptide contributed, on average, <1% of the total circulating GDF15 (Median Percentage maternal GDF15: 0.60% [Q1, Q3: 0.12, 2.25]) (**Figure 2B-D**). The maternal fraction of GDF15 appeared to increase in some pregnancies between the first and second trimester but declined in later pregnancy (**Extended Data Figure 2A**) as circulating concentrations of total GDF15 rise (**Extended Data Figure 2B**). To confirm that antenatal circulating GDF15 was near-exclusively of fetal origin, we repeated these experiments using samples from maternal plasma where the fetus was heterozygous at the H202D position, and the mother was homozygous for the reference allele. Surprisingly the D-peptide, which is produced only by the fetus in this cohort, constituted greater than half of the total circulating GDF15 (Mean percentage D-peptide: 62.6%, 95%CI[59.1, 66.0], P=6.80 x 10^−6^, one-sample T-Test) – implying that it was present in excess of what would be expected even if all circulating GDF15 was fetal in origin (**Figure 2E**). This was not attributable to assay bias (**Extended Data Figure 2C**). These data suggest that the D-peptide may be preferentially secreted or may have a prolonged half-life in the circulation.

**Figure 2.**
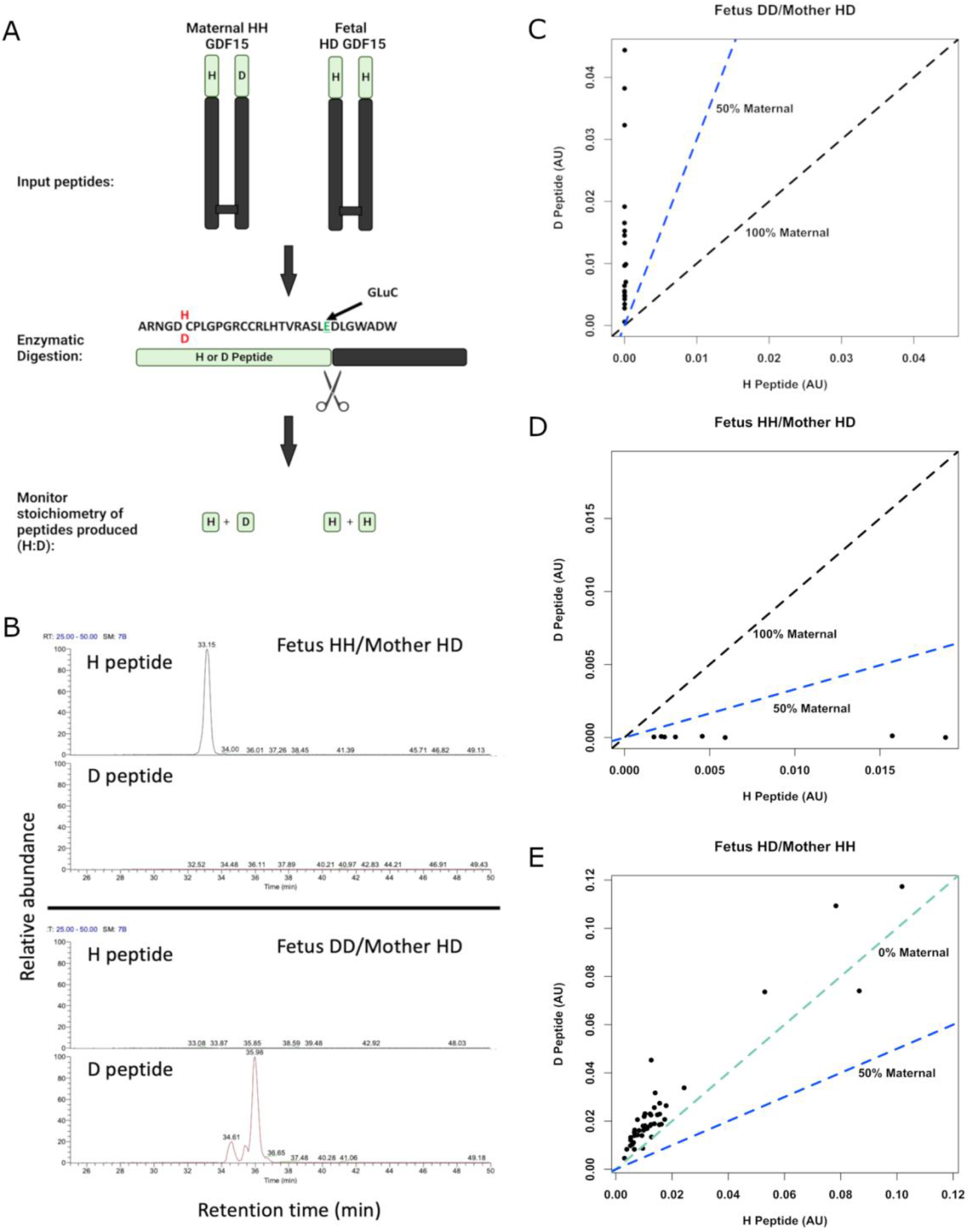
Circulating GDF15 in human pregnancy is predominantly of fetal origin. **A**: Schema of experimental design. The GDF15 dimer for maternal and fetal GDF15 is extracted and then digested with the endopeptidase GluC, cutting the N-terminal region into two distinct peptides with glutamic acid C-termini. The stoichiometry of the H and D peptides can then be monitored using LC-MS/MS to determine the relative levels of maternal or fetal derived GDF15 in the maternal circulation. **B**: Representative LC-MS retention time of H and D peptides from maternal plasma where the mother is heterozygous at H202D and the fetus is homozygous for the H or D allele as indicated. **C-E:** Scatter plots of the relative quantitation of H peptide vs the D peptide in plasma from pregnancies with the indicated genotypes. The dashed coloured lines in **(C-E)** indicate the expected relationships between the H and D peptides for the given circulatory origins of GDF15. **C:** N=20 samples from 5 pregnancies, **D**: N=8 samples from 2 pregnancies, **E**: N = 47 samples from 12 pregnancies.

### A rare genetic variant in GDF15 that predisposes to Hyperemesis Gravidarum results in impairment of GDF15 secretion and reduced circulating GDF15 in the non-pregnant state

Fejzo et al have previously reported that women heterozygous for the C211G mutation in GDF15 have at least a 10-fold increased risk of developing HG [10]. Cysteine 211 is one of the key conserved cysteine residues involved in intrachain di-sulphide bonding of GDF15 and its absence would be predicted to be highly damaging [12]. Supporting this, when we transiently transfected a construct encoding GDF15 with a glycine at position 211 into HEK 293T cells it was highly expressed but, unlike wild-type, the mature peptide was not secreted and the unprocessed pro-peptide was completely retained intracellularly (**Figure 3A, Extended Data Figure 3A**). GDF15 is secreted as a homodimer, so we wished to test whether the mutant form might interfere with the secretion of wild-type GDF15. We differentially tagged mutant and wild-type forms of GDF15 and demonstrated a clear reduction in the secretion of wild-type GDF15 when it was co-expressed with 211G (**Figure 3B, Extended Data Figure 3B-C**).

**Figure 3.**
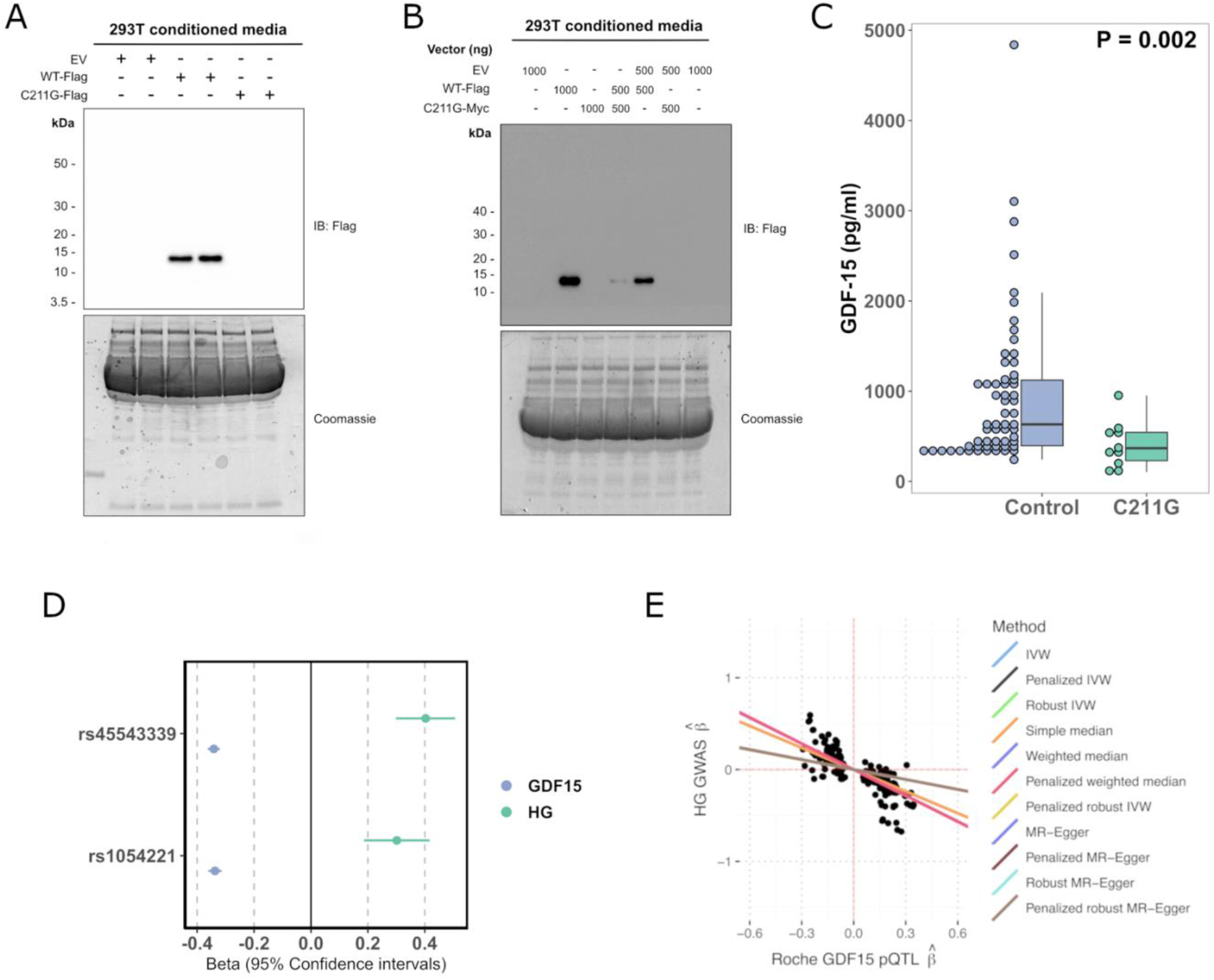
Rare and common hyperemesis gravidarum risk variants lower circulating GDF15 in the non-pregnant state. **A:** We studied the secretion of a rare hyperemesis gravidarum risk variant GDF15 C211G. C211G impairs the secretion of GDF15 as determined by western blotting of cell culture medium of cells expressing Flag-tagged wild-type GDF15 (WT-Flag) or GDF15 C211G (C211G-Flag). **B:** GDF15 C211G impairs the secretion of wild-type GDF15 in a dominant negative manner as co-expression of the mutant inhibited secretion of wild-type GDF15 from 293T cells co-transfected with different amounts (shown in nanograms) of wild-type WT-Flag and Myc-tagged GDF15 C211G (C211G-Myc), as indicated. For A and B representative images from 3 independent experiments are presented. EV represents transfection with the empty plasmid backbone. **C:** Dot and box plots showing GDF15 levels measured using the Ansh Total GDF15 assay in carriers of GDF15 C211G variant (N=10) identified in an exome-sequencing study of a Croatian population and age and sex matched controls (N=60) derived from the same study. The P-value is from a linear regression model of natural log transformed GDF15 ∼ C211G status. **D:** Forest plot illustrating the effect (standardised betas) of previously described HG risk SNPS on circulating GDF15 measured in 18,184 participants in the Generation Scotland Study. The effect estimates for the rs1054221 variant presented are from an analysis conditioned on the the lead HG variant rs45543339. The beta estimates for GDF15 represent the effect of the HG risk allele on circulating GDF15 in standard deviations. The beta estimates for HG (Hyperemesis Gravidarum) represent the effect of the SNP on risk of HG in log-odds. **E:** Scatterplot of HG GWAS effect estimates (ie log-odds) vs Roche-based GDF15 pQTL effect estimates derived from cis-Mendelian randomization at the *GDF15* locus. MR was performed using m=259 SNPs with genome-wide evidence of pQTL effects on GDF15 levels within 1Mb *GDF15* locus and adjusted using LD estimates from UK Biobank WGS individuals (n=138335; see Methods). Causal effect estimates obtained using LD-aware MR and reflected as regression lines.

To determine the effect of the C211G variant on circulating GDF15, we identified a Croatian cohort [13] in which exome sequencing had identified 11/2872 C211G heterozygotes (Minor Allele Frequency ∼ 0.002). Levels of circulating GDF15 (measured by an in-house MSD assay using the Ansh Lab Total GDF15 antibodies) in C211G heterozygotes (none of whom were known to be pregnant) were reduced by more than 50% compared to age and sex matched controls from the same population (**Figure 3C, Supplementary Table 7**).

To clarify the interaction between maternal and fetal carriage of the C211G variant, we identified 17 offspring of six women previously found to be heterozygous for C211G [10]. The mothers had HG in 10/10 pregnancies where the fetus was homozygous for wild-type at position C211. Conversely, HG was reported in only 4/7 pregnancies where the fetus was heterozygous for C211G (**Supplementary Table 8**), suggesting that maternal carriage of the C211G variant confers HG risk and that this risk may be moderated when the variant is also carried by the fetus.

### Common genetic variants predisposing to HG are associated with lower circulating GDF15 levels in the non-pregnant state

Common genetic variants in and around the *GDF15* gene have been reported to have the strongest genome wide association with HG. We studied two single nucleotide variants at the *GDF15* locus which are independently associated with HG [9] and examined their association with GDF15 levels (measured by Roche Elecys) in 18,184 people from the Generation Scotland Study [14]. Consistent with the effects of the rare C211G variant, both HG risk alleles are associated with lower GDF15 in the non-pregnant state (rs45543339: β = -0.34, 95%CI[-0.36, -0.32], rs1054221 conditioned on lead signal: β = -0.34, 95%CI[-0.36, -0.31] **Figure 3D**).

To systematically test for a causal relationship between circulating GDF15 in the non-pregnant state and HG risk, we performed LD-aware Mendelian randomization (MR) analysis using cis-pQTLs (P < 5x10^−8^) identified in a genome wide association study of circulating GDF15 (measured by Roche Elecys) in the Generation Scotland Study (N=18,184). Overall, we observed that increased circulating GDF15 in the non-pregnant state reduced HG risk (IVW MR; OR=0.70 per SD increase in GDF15 95%CI[0.65-0.76], P = 6.98 x 10^−17^) (**Figure 3E, Supplementary Table 9**). These results were robust to the choice of LD reference panel and MR approach (**Extended Data Figure 4, Supplementary Table 9-10**). While we have previously demonstrated that the Roche Elecsys assay is not affected by the common protein altering variant H202D (rs1058587) [11], we wished to exclude any possibility that small biases in detection related to this variant could explain our findings. Therefore, we repeated our analysis after conditioning on this variant producing similar results (**Extended Data Figure 5, Supplementary Table 11**).

**Figure 4.**
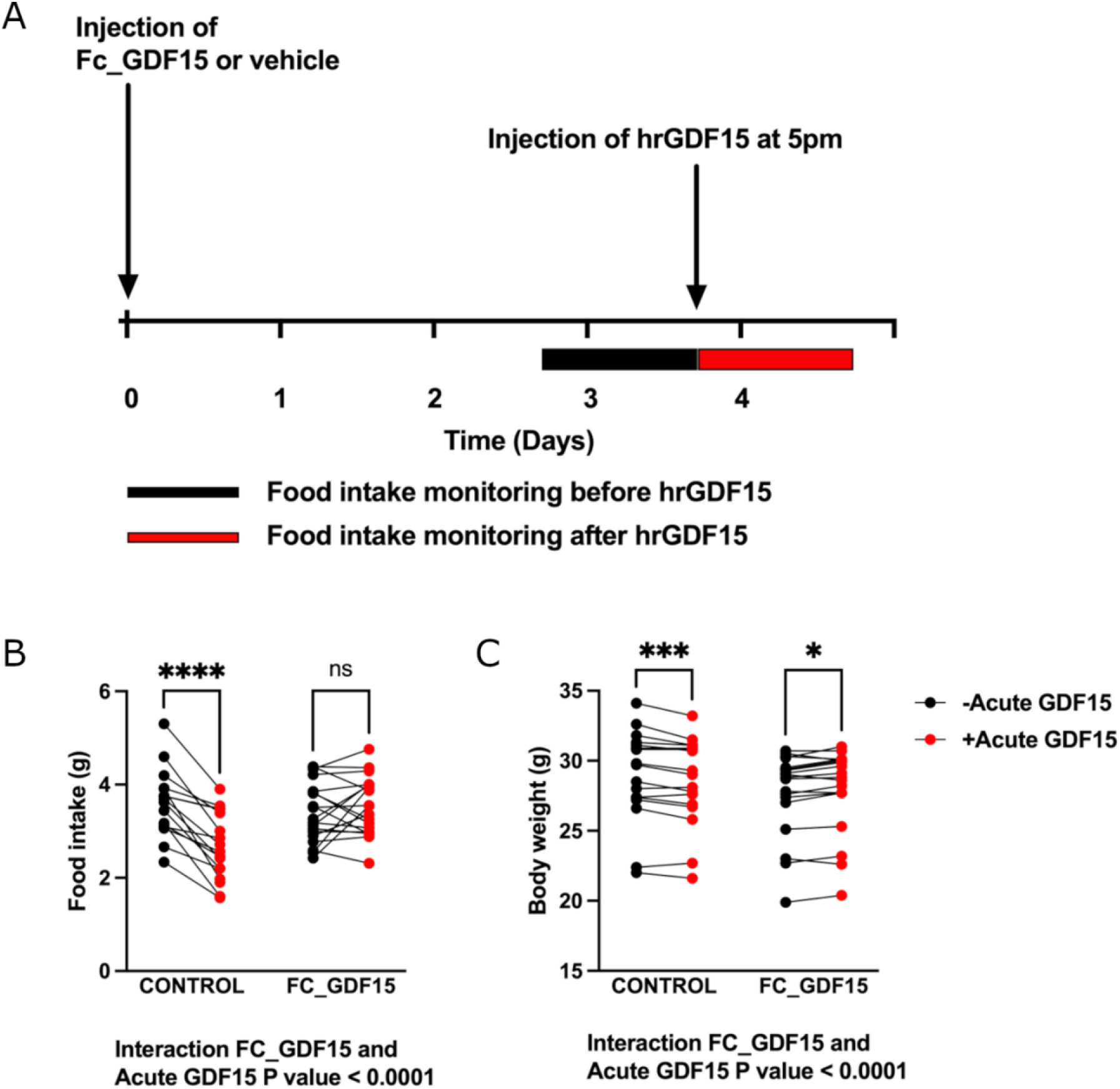
Treatment with long acting GDF15 influences the response to the anorectic actions of acute GDF15 treatment in mice. **A**: Schema of the experimental paradigm. Single housed, adult male and female C57Bl/6J mice were injected with 0.01mg/kg of Fc-GDF15-15 fusion protein (Fc_GDF15) or vehicle control (PBS). Food intake was measured overnight (from 5pm to 9am) before (black bar) and after treatment (red bar) with standard, short acting human recombinant GDF-15 (hrGDF15 0.1mg/kg). Body weight was measured at the same timepoints. **B**: Food intake recorded overnight (5pm-9am) the day before (black dots) and after an acute bolus of hrGDF15 (red dots) in mice with and without pre-treatment with Fc_GDF15. **C:** Body weight at 9am the day before (black dots) and 9 am the day after (red dots) an acute bolus of hrGDF15 in mice with and without pre-treatment with Fc_GDF15. N=17 (12 male, 5 female) in Control and 19 in FC_GDF15 group (13 male, 6 female). Data were analysed using mixed-effect analysis, post-hoc testing comparing before and after acute GDF-15 treatment was undertaken with the Sidak test to correct for multiple testing. *P<0.05, ***P<0.001, ****P<0.0001, ns = non-significant, P>0.05.

Finally, we used statistical colocalization implemented in *coloc*, a complementary approach to MR which can be used to assess the probability that a genetic signal is shared between an outcome of interest and an intermediate molecular trait, in this case HG and circulating GDF15, respectively. We observed two colocalizing signals at the *GDF15* locus (rs45543339 and rs1227731; PPH4 >0.99, **Supplementary Table 12**), which correspond to the two independent signals presented in Figure 3D, where both HG risk-raising alleles are associated with reduced GDF15 in the non-pregnant state.

Thus, from studies of both rare and common genetic variants in GDF15, it appears that higher circulating levels of the hormone in the non-pregnant state are associated with protection from HG.

### Prior exposure to GDF15 influences responses to an acute bolus of GDF15

To test the hypothesis that prior levels of exposure could influence acute responses to GDF15 we administered a long-acting form of GDF15 (FC human GDF15, 0.01mg/kg, [15]) to wild-type mice (**Figure 4A**). Pre-treatment with FC human GDF15 resulted in a mean concentration of 4773 ± 440pg/ml 3 days after the injection, which corresponds to ∼47-fold increase compared to basal circulating levels of mouse GDF15. This results in transient suppression of food intake for one day after injection relative to untreated controls (**Extended Data Figure 6**). 3 days after treatment with FC human GDF15, mice were then given an acute bolus of human recombinant GDF15 (0.1mg/kg), which typically elevates GDF15 to >20,000 pg/ml 1 hour after injection [16], and its effects on food intake and body weight were measured. Mice previously receiving a vehicle control injection showed the expected major reduction in food intake in response to a bolus of GDF15 (**Figure 4B**) and lost weight (**Figure 4C**). In contrast, mice previously exposed to GDF15 had a markedly blunted acute response (**Figure 4B-C**), supporting the notion that elevated antecedent levels of GDF15 can influence the subsequent action of an acute rise in circulating GDF15.

### Pre-pregnancy exposure to high levels of GDF15 protect against the development of nausea and vomiting

Some chronic medical conditions are characterised by life-long elevations of GDF15. Our hypothesis predicted that such exposure might reduce the risk of developing nausea and vomiting when those individuals become pregnant. Beta thalassemia is a genetic disorder affecting red blood cells where extremely high levels of GDF15, found throughout life [17, 18], are thought to come from the expanded mass of stressed erythroblasts. Though fertility is impaired in this disease, some women, particularly those with thalassemia intermedia, do become pregnant. We conducted a survey (**see Supplemental material**) of women with beta-thalassaemia who had undergone at least one pregnancy which had resulted in a live birth and compared the results with ethnically- and age-matched women who did not have thalassemia. There was a striking reduction in symptoms of NVP in the women with thalassemia: only ∼5% of women with thalassemia reported any nausea or vomiting compared to >60% of the controls (P<0.01) (**Supplementary Table 13**).

### Summary and conclusions

Despite the fact that nausea and vomiting are symptoms which occur in most human pregnancies, are commonly disabling and, when severe, can be life-threatening, their aetiology and pathogenesis have remained poorly understood. Here we present evidence that the severity of nausea and vomiting of pregnancy is the result of the interaction of fetal derived GDF15 and the mother’s sensitivity to this peptide, which is substantially determined by her prior exposure to the hormone.

Using immunoassays that are not confounded by the common H202D variant we showed that levels of GDF15 in the maternal circulation in the late first trimester are significantly increased in women with HG compared to those without severe nausea and vomiting and also in an independent cohort of pregnant women who, early in the second trimester, reported vomiting in pregnancy compared to those reporting no nausea or vomiting. We can now conclude with confidence that higher circulating levels of GDF15 in maternal blood are associated with an increased risk of HG. However, as there is considerable overlap in levels between HG cases and controls, GDF15 concentrations alone cannot be used as a diagnostic tool to differentiate HG from other causes of vomiting in a pregnant woman.

We employed mass spectrometry applied to genetically discordant mother/offspring pairs and identified the fetal genome as the dominant source of the circulating GDF15 in maternal blood. This finding is consistent with previous reports of extremely high levels of *GDF15* gene expression in, and protein secretion from, human trophoblast [5]. A caveat to this observation is that these studies were undertaken in healthy pregnancies, and it is conceivable that, in women with established HG, stressed maternal tissues may, in theory, make an additional contribution to the circulating pool.

The rare coding variant GDF15 C211G has been reported to greatly increase the risk of HG [10]. We report that this mutation is associated with markedly lower circulating levels in the non-pregnant state attributable to the deleterious effects of the mutation on secretion of mature GDF15, including any wild-type partner present in heterodimers. We also demonstrate that common HG risk conferring variants are associated with lower circulating levels of GDF15 in the non-pregnant state. Conversely, high levels of GDF15 preceding pregnancy, as are found in thalassemia, appear to be strongly protective against the development of NVP. This finding is consistent with studies which report that pre-pregnancy cigarette smoking (a behaviour associated with elevated GDF15 [19], reduces the risk of HG [20].

Agonist induced desensitisation is a feature of many hormone-receptor systems and here we show that this occurs in the case of GDF15, and its receptor GFRAL-Ret. Mice exposed to mildly supraphysiological doses of GDF15 for 3 days show markedly attenuated food intake and body weight responses to an acute bolus of GDF15. The tendency for the GDF15/GFRAL-Ret system to exhibit some degree of ligand induced desensitisation provides a plausible explanation for the effects of pre-pregnancy GDF15 exposure on the risk of NVP and HG developing in the face of the acute increase in circulating GDF15 which begins in early pregnancy.

We report that levels of GDF15 are higher in pregnant women with NVP and HG than in those without those symptoms and have also shown that the feto-placental unit is the major source of that GDF15 in maternal blood. Mothers with HG are enriched in GDF15 variants which are associated with lower GDF15 in the non-pregnant state and will transmit ∼50% of those alleles to their offspring, in whom they might be expected to lower GDF15 levels. How can those observations be reconciled? Firstly, it is possible that variants that affect the expression of GDF15 do so differentially in adult tissue vs placenta. Secondly, there are factors beyond the GDF15 gene which may influence GDF15 production by the feto-placental unit. For example, female fetal sex, the presence of twins, or the presence of invasive trophoblastic disease are all associated with increased HG risk [21, 22] and, at least in the case of female fetuses, increased GDF15 levels in pregnancy [23]. In the case of the C211G HG risk variant, we found suggestive evidence for an interaction of maternal and fetal GDF15 genotype with fetal carriage of this variant apparently moderating the maternal effect. Thus, HG occurred in 10 out of 10 pregnancies where the mother was a C211G heterozygote (presumably with low pre-pregnancy levels of circulating GDF15) and was carrying a wild-type fetus, but only in 4 out of 7 pregnancies where the fetus was heterozygous for the mutation. Future studies examining the relationship between maternal and fetal genotype, HG symptoms and maternal GDF15 levels in large sample sizes should provide further clarification.

Our findings have obvious implications for the prevention and treatment of HG. The acute rise in GDF15 which accompanies normal pregnancy is, we would argue, likely to be necessary, if not sufficient, for the causation of HG. The corollary of this is that blocking GDF15 action in the pregnant mother should be a highly effective therapy for women suffering from HG. We make this argument based on a number of observations. Firstly, the administration of acute bolus of GDF15 to humans, resulting in levels similar to that seen in pregnancy, frequently results in nausea and vomiting [24]. Secondly, in non-human primates, blocking GDF15 is highly effective in reducing vomiting resulting from the administration of drugs like cis platinum which cause an acute increase in GDF15. Thirdly, human genetic variants, common and rare, point to the GDF15 system as a major susceptibility locus for human HG. Fourthly, the striking reduction in frequency of NVP in women with thalassemia, a condition of markedly increased pre-pregnancy levels of GDF15, suggests that GDF15 plays a key role in the causation of these symptoms in pregnancy. The fact that high GDF15 levels in the non-pregnant state appears to be protective against the development of NVP and HG suggests that strategies which safely increase circulating GDF15 levels prior to pregnancy may be useful in the prevention of these conditions.

Since the tragedy of thalidomide [25], concerns about safety have understandably been very prominent in discussions of novel treatments for HG, particularly any that would cross the placenta and carry a risk of teratogenesis. For other disease indications, antibodies have been engineered to minimise their transplacental passage, and have been widely used [26], so this should be a possible route to safe blockade of GDF15 signalling. There are reasons to think that highly specific blockade of GDF15 signalling through its receptor GFRAL is likely to be safe, even if such an antagonist did gain access to the fetus. GDF15 appears to act highly specifically through GFRAL, which is only expressed in the hindbrain. Mice lacking GDF15 or GFRAL develop normally and remain largely healthy throughout life.

GDF15 appears to have evolved primarily as a signal to confer information about a range of somatic stresses (such as produced by toxins) to the brain in order to reduce continuing exposure to those stresses at the time of exposure and in the future, through promoting avoidance behaviour [27]. The placentae of certain higher mammals, including primates, have evolved to produce large amounts of GDF15 from early pregnancy, a phenomenon which likely explains the very common occurrence of nausea and vomiting in pregnant women [5]. Sherman and Flaxman [28] suggested that the evolutionary basis for this likely lay in the protection of both mother and fetus from food-borne illness and toxins, particularly important at a time when the fetus is most susceptible to teratogens and the immunosuppressed state of early pregnancy made mothers susceptible to infections. The energy needs of the growing fetus may outweigh the risks as the pregnancy progresses resulting in the selection against persistence of NVP beyond the 1^st^ trimester in normal pregnancies. The phenomenon of ligand induced desensitisation, which we have demonstrated to occur with GDF15 may explain the natural tendency for the severity of NVP to wane as pregnancy progresses.

In conclusion, our findings place GDF15 at the mechanistic heart of NVP and HG and clearly point the way to strategies for its treatment and prevention.

## Methods

### Cambridge Baby Growth Study

The CBGS is a prospective, longitudinal cohort study originally recruiting 2,229 pregnant women from the Rosie Maternity Hospital, Cambridge between April 2001 and March 2009 [7]. This analysis was performed using a nested case-control format from those that returned filled-in pregnancy questionnaires, including questions about nausea and vomiting in pregnancy, and who provided a blood sample taken between 12 and 18 weeks of pregnancy. The cases were women who reported vomiting in pregnancy, and the controls were women who reported neither nausea nor vomiting in pregnancy. The samples for GDF15 measurement were chosen according to availability. The statistical analysis was performed using linear regression (and natural log-transformed GDF15 concentrations so that the residuals were normally distributed), either unadjusted or adjusted for potential confounders such as gestational age at sampling and body mass index. Ethical approval for the Cambridge Baby Growth Study was granted by the Cambridge Local Research Ethics Committee, Cambridge University Hospitals NHS Foundation Trust, Cambridge, U.K. (00/325).

### HG Study

The HG Study is a case-control study of women recruited from the Rosie Maternity Hospital, Cambridge and North West Anglia (NWA) NHS Foundation Trust at Peterborough City Hospital, between 2018 and 2021. The 72 cases were pregnant women admitted to hospital for rehydration due to hyperemesis gravidarum. The 182 controls were pregnant women admitted to hospital in the same pregnancy timeframe as the cases, but for other reasons (e.g. termination of pregnancy or uterine bleeding). Blood samples were collected around week 9 of pregnancy, and a nausea/vomiting score was calculated by asking the women for their current and worst nausea and vomiting ratings out of ten. The samples for GDF15 measurement were chosen to maximise the difference in the nausea/vomiting scores. The statistical analysis was performed using linear regression (and natural log-transformed GDF15 concentrations so that the residuals were normally distributed), either unadjusted or adjusted for potential confounders such as gestational age at sampling. Ethical approval was granted by the National Research Ethics Service Committee - East of England, Norfolk, U.K. (14/EE/1247). All procedures followed were in accordance with both institutional and international guidelines. Written informed consent was obtained from all women.

### C211G carriers and Controls in the CROATIA-Korcula Study

The CROATIA-Korcula study sampled 2926 Croatians from the Adriatic island of Korcula, between the ages of 18 and 98. The fieldwork was performed from 2007-2014. Ethical approval was given for recruitment of all participants by ethics committees in both Scotland and Croatia. All volunteers gave informed consent before participation. Carriers of GDF15 C211G variant with available serum samples were identified from the exome-sequence of samples from the CROATIA-Korcula study and were paired with age and sex matched controls from the same cohort.

### Common genetic variation, circulating GDF15 and risk of hyperemesis gravidarum

#### 23andMe HG GWAS data

We obtained 23andMe, Inc (23andMe) GWAS summary statistics of HG from ref [9]. Briefly, 23andMe GWAS research participants provided answers to morning sickness-related questions. All research participants provided informed consent and volunteered to participate in the research online, under a protocol approved by the external AAHRPP-accredited IRB, Ethical & Independent (E&I) Review Services. As of 2022, E&I Review Services is part of Salus IRB (https://www.versiticlinicaltrials.org/salusirb). HG status was defined as 1306 research participants who reported via an online survey that they received IV therapy for NVP and 15,756 participants who reported no NVP served as controls. For additional details refer to ref [9].

#### GDF15 pQTL data and quality control

Generation Scotland is a family and population-based study consisting of 23,690 participants recruited via general medical practices across Scotland between 2006 and 2011. The recruitment protocol and sample characteristics are described in detail elsewhere [29, 30]. Ethical approval for the Generation Scotland study was obtained from the Tayside Committee on Medical Research Ethics (on behalf of the National Health Service).

The GWAS analysis used BOLT-LMM in order to adjust for population structure and relatedness between individuals [31] in a linear mixed model analysis of Generation Scotland participants with available GDF15 data and Haplotype Reference Consortium reference panel release 1.1 [32, 33] imputed genotype information (18184 individuals). Age, sex and first 20PCs were included as covariates. Serum GDF15 concentrations were subject to rank-based inverse normal transformation prior to analysis. Associations were considered significant when P ≤ 5 × 10—8.

#### Conditional GWAS analyses

To assess the extent to which signals beyond lead (or focal) SNPs contribute to either HG risk (or GDF15 levels), we performed conditional analyses using GWAS summary data and estimates of LD derived from the regression of summary statistics model (i.e. RSS) [34]. Briefly, given estimated effect-sizes β(e.g., log-odds or linear effects) at *m* non-leading SNPs, corresponding *m* standard errors *s, m* × *m* LD matrix *V*, and *m* × 1 vector *v* of LD estimates with the lead SNP, we can compute residual effect-sizes *β*\* as,

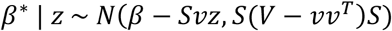

where *S* = *diag(s)* is the *m* × *m* diagonal matrix of standard errors, *z* = *b*/*se*(*b*) is the association statistic at the lead (or focal) SNP, and *N*(·,·) corresponds to the multivariate normal distribution. The conditional estimates correspond to the mean of the above distribution and standard error proportional to the diagonal of the covariance.

To compute conditional effect-size estimates for circulating GDF15 levels and separately for HG risk, we used the above model focusing on m=310 harmonized variants and LD estimates from WGS data in European-ancestry individuals in the UK Biobank cohort (see below).

#### Mendelian randomization analyses

To perform Mendelian Randomization between circulating GDF15 levels with HG risk, we harmonized Roche-based GDF15 pQTL, GWAS, and LD reference panels to obtain valid estimates. First, we restricted analysis to variants associated with GDF15 levels at a genome-wide significant threshold (p<5e-8) ±500Kb around the transcription start site. Next, we harmonized GDF15 pQTL significant results with 23andMe HG GWAS association statistics to match for consistent reference and alternative alleles, which resulted in m=311 variants. We excluded any variants whose reference and alternative alleles may be ambiguous (e.g., G/C, A/T), except for previously referenced risk alleles (e.g., rs1058587). To account for linkage between GDF15-associated variants, we estimated LD using WGS data from European-ancestry individuals in the UK Biobank (UKBB) cohort (n=138355) as well as WGS data from European-ancestry individuals in the 1000G study (n=489). To derive LD estimates in UKBB the publicly available whole-genome sequencing (WGS) data from European participants in UKBB (n = 138335) was used for the determination of linkage disequilibrium at the GDF15 locus. 5259 WGS variants were extracted ±500KB from chr19:18388612:C:G (GRCh38) and the Pearson’s R was determined using PLINK v1.90b6.26/Swiss Army knife App via the UKB Research Access Platform, with the following parameters ‘--ld-window-r2 0 --ld-window 10000 --keep-allele-order --snp chr19:18388612:C:G --window 1000’. All work using the UKBB resource was conducted using application numbers: 9905 and 32974.

Harmonizing our association data with LD estimates resulted in m=259 variants for UK Biobank data and m=310 variants when using 1000G data. Lastly, we performed Kriging analysis using the R package susieR (https://cran.r-project.org/web/packages/susieR/) to ensure no variants were mislabelled between reference LD and association study results. Finally, to perform Mendelian Randomization, we used the R package MendelianRandomization (https://cran.r-project.org/web/packages/MendelianRandomization/index.html).

#### Colocalization analyses

To perform colocalization analysis between genetic variants underlying circulating GDF15 levels and HG risk, we performed the same harmonization strategy as the LD-aware Mendelian Randomization analysis in Roche-based GDF15 pQTL data and 23andMe GWAS results. However, rather than limit analyses to variants with genome-wide significance for pQTL effects, we selected all variants represented in LD estimated from UK Biobank WGS data, which resulted in m=2,297 variants. We performed multi-causal SNP colocalization using the R package coloc (https://cran.r-project.org/web/packages/coloc/index.html), which tests for colocalization across SNPs identified within credible sets, to better reflect linkage patterns.

### Prevalence of nausea and vomiting in pregnancy in thalassaemia

We conducted a survey to compare the prevalence of NVP among women with beta-thalassaemia and ethnically- and age-matched non-thalassaemia healthy women at the Colombo North Teaching Hospital, Ragama, Sri Lanka from 01 June to 31 August 2022. All female patients with beta-thalassaemia with at least a single child attending for regular blood transfusions and thalassaemia follow-up during the study period were recruited. An equal number of ethnically- and age-matched non-thalassaemia healthy females with at least a single child attending the general paediatric clinic of the same hospital with their children during the study period were recruited as controls. Specifically, we recruited the eligible ethnically- and age-matched non-thalassaemia control attending the clinic on the same day immediately after recruiting a beta-thalassaemia patient. Informed written consent was obtained from all study participants before recruitment. Data on nausea, vomiting and loss of appetite during pregnancy were gathered using an interviewer-administered questionnaire **(see Supplemental material)**. The prevalence of nausea, vomiting and loss of appetite during pregnancy of beta-thalassaemia patients and non-thalassaemia women were compared using logistic regression after adjusting for parity, number of children and time since index pregnancy. The study was approved by the Ethics Review Committee of University of Kelaniya, Sri Lanka (Ref: P/228/11/2019).

### Maternal NVP levels and offspring genotype

Carriers of rs372120002 (C211G) were identified in a previous whole-exome sequencing study of Hyperemesis Gravidarum [10]. Eleven carriers of rs372120002 (C211G) and their children were invited to participate in the offspring study, among which 6 carrier mothers and 17 children agreed to participate. Participating mothers filled out a survey on NVP/HG during each of their pregnancies which included whether they had HG, were treated with antiemetic medication(s) and intravenous fluids, had an emergency room visit and/or hospitalization for HG, and when their symptoms resolved. Cheek swab samples were collected from children using DNA Genotek cheek swab kits (OCD-100, OC-175, Oragene, Ottawa, Canada), and DNA was extracted according to the manufacturer’s recommendations. PCR of rs372120002 was performed using standard methods with forward primer CAGCTCAGCCTTGCAAGAC and reverse primer GGATTGTAGCTGGCGGGC, annealing temperature at 60 °C, and the PCR product was sequenced by Azenta, Life Sciences (Chelmsford, MA). Genotypes were called using 4Peaks app to view DNA trace files. The study was approved by the USC Institutional Review Board.

### GDF15 immunoassays

Total GDF15 levels in the CBG cohort were measured using a 3-step plate ELISA (Ansh AL-1014-r) which was validated to be able to recognise H and D containing variants at position 202 (position 6 of the mature peptide) of GDF15 with comparable affinity (**Supplementary Table 1**). The calibrators, kit controls, in-house sample pool controls (diluted 1:15 in Sample Diluent) and samples (diluted 1:15 in Sample Diluent) were added to the antibody coated microtiter plate and incubated. Following a wash step the biotinylated detector antibody was added and incubated. Following a second wash step streptavidin horse radish peroxidase conjugate solution was added and incubated. Following a third wash step substrate solution (TMB) was added and incubated followed by an acidic stop solution. The measured absorbance at 450nm corrected at 630nm is directly proportional to the GDF15 concentration. The calibrator supplied with the kit by Ansh Labs is traceable to recombinant human GDF-15 from R&D Systems (Biotechne, USA). Ansh Lab ELISA Total GDF15 between batch imprecision kit controls 7.7% at 173.2 pg/ml, 5.1% at 480.0 pg/ml and in-house sample pool controls 4.9% at 397.5pg/ml, 3.7% at 1022.5 pg/ml

GDF15 measurements in both the HG vs control study and in Generation Scotland were measured on a Cobas e411 analyser (Roche Diagnostics, Basel, Switzerland) using the manufacturer’s reagents and quality control material. Coefficient of variation for GDF15 was 3.8% for the low control (at 1,556 pg/mL) and 3.4% for the high control (at 7,804 pg/mL). The limit of detection (LoD) of the GDF15 assay is set to 400 pg/mL by the manufacturer, and the upper limit of the measuring range was 20,000 pg/mL. As previously reported [14] for the Generation Scotland study, for continuous analysis, samples below the limit of detection were reported as 200 pg/mL and samples above the measuring range as 25,000 pg/mL. For the HG vs Control pregnancy study, samples were diluted 1 in 5 with assay buffer before measurement because of the known very high levels in pregnancy. To examine the effect of the H202D variant on GDF15 immunoreactivity we determined the recovery of synthetic peptides produced as previously described [11].

Total GDF15 levels in the Croatia-Korcula study were measured using an in-house developed assay on the Meso Scale Discovery (MSD) platform using two monoclonal antibodies from Ansh Labs which have been described as being able to recognise H and D containing variants at position 202 (position 6 of the mature peptide) of GDF15 with comparable affinity. The calibrators, in-house sample pool controls and samples were added to the monoclonal antibody coated MSD plate and incubated. Following a wash step the biotinylated detector monoclonal antibody diluted in MSD Diluent 100 was added and incubated. Following a second wash step Sulpho-TAG labelled Streptavidin (MSD) diluted in MSD Diluent 100 was added and incubated. Following a third wash step MSD read-buffer was added to all the wells and the plate was immediately read on the MSD s600 plate reader. Luminescence intensities for the standards were used to generate a standard curve using MSD’s Workbench software package and were directly proportional to the GDF15 concentration. The calibrator is recombinant human GDF15 from R&D Systems (Bio-Techne, USA). MSD Ansh antibody Total GDF15 between batch imprecision based on in-house sample pool controls 10.2% at 552.4 pg/ml, 11.7% at 1518.6 pg/ml, 11.7% at 7036.1pg/ml.

### Identification of mother/fetus pairs discordant for H202D

Mother-offspring pairs not fully concordant for genotype at the H202D site in GDF15 were identified by first genotyping the offspring using placental RNA sequencing data from the Pregnancy Outcome Prediction (POP) study cohort [6]. We used the GATK pipeline [35] to identify SNPs (Single Nucleotide Polymorphisms) from the RNA-Seq alignment data (i.e., BAM files). Briefly, the pipeline comprises the following steps: (1) marking duplicate reads using ‘markDuplicate’ of Picard (https://broadinstitute.github.io/picard/), (2) splitting reads that contain ‘N’s in their CIGAR string using ‘splitNRead’ of GATK (subsequent submodules from GATK hereafter), (3) realignment of reads around the indel using ‘IndelRealigner’, (4) recalibrating base quality using ‘BaseRecalibrator’, and (5) calling the variants using ‘HaplotypeCaller’. Homozygous alternative alleles and their read counts were parsed directly from the VCF files generated by the previous step, 5). As homozygous reference alleles are not called by ‘HaplotypeCaller’, we used ‘mpileup’ command of samtools and bcftools to detect the read counts from the BAM files generated by the previous step. For heterozygous SNPs, we counted reads by the reference and alternative bases using ‘ASEReadCounter’. Fetal genotype was confirmed using umbilical cord DNA and the maternal genotype determined using the TaqMan™ SNP Genotyping Assay to rs1058587 (Applied Biosystems) according to the manufacturer’s instructions.

### Mass spectrometry studies

Anti-human GDF15 capture antibody (R&D systems) was coupled to tosyl-activated M-280 paramagnetic dynabeads (ThermoFisher Scientific) using the standard supplied protocol. Plasma from each individual (50 µL) was diluted with 150 µL of Buffer E and 5 µL of magnetic beads at 20 mg/mL was added. Samples were mixed at 850 rpm for 1 hour at room temperature on a 96 well MixMate plate mixer (Eppendorf). The beads were concentrated using a magnet and the supernatants removed. The beads were washed twice in 200 µL of buffer E. A final wash with 200 µL of 50 mM ammonium bicarbonate was performed and the supernatant removed. Disulphide bonds were reduced with 75 µL of 10mM DTT in 50 mM ammonium bicarbonate over a 60-minute incubation at 60°C, before alkylation with 20 µL of 100 mM iodoacetamide in 50 mM ammonium bicarbonate in the dark for 30 minutes at room temperature. To digest the polypeptide 10 µL of Glu C enzyme (Worthington) at 100 µg/mL was added and the samples digested overnight at 37 °C. The digestion was stopped by the addition of 20 µL of 1% formic acid in water.

The plasma samples collected from mothers with homozygous fetuses were analysed on a ThermoFisher Q-Exactive Plus Orbitrap using nanoflow analysis with an Ultimate 3000 LC system. Peptides monitored were ARNGD**<u>H</u>**CPLGPGRCCRLHTVRASLE and ARNGD**<u>D</u>**CPLGPGRCCRLHTVRASLE corresponding to the H-peptide and D-peptide (mutant) respectively. Additionally, a GluC derived peptide was monitored from the murine anti-human GDF15 antibody as a surrogate internal standard with which to generate a peak area ratio for comparing relative peptide levels. This peptide was FKCKVNNKDLPSPIE from the heavy chain. A parallel reaction monitoring method was developed for the GDF15 peptides targeting the [M+5H]^5+^ ion, which corresponded to 579.08 and 574.67 *m/z* for the H and D peptide respectively. The Product ions corresponding to the same y18 ion for each peptide (693.6950, 694.0285 and 694.3614 *m/z*) were summed for quantitative analysis in the Quan Browser program (ThermoFisher). The plasma samples from mothers with heterozygous fetuses were analysed on an M-Class LC system (Waters), linked to a Xevo TQ-XS triple quadrupole mass spectrometer (Waters) with an IonKey interface. SRM transitions used for these peptides were 579.24/693.89 and 579.24/747.25 for the H peptide, 574.82/693.89 and 574.82/623.73 (the first SRM transition for each peptide was used as the quantifier transition) as well as 545.27/682.34 and 545.27/926.37 which targeted the peptide ATHKTSTSPIVKSFNRNEC from the C-terminus of the murine antibody kappa light chain. In both experiments, peptides from the GDF15 protein were expressed as peak area ratios relative to the murine antibody peptide.

#### Estimating total and maternal derived GDF15 by mass spectrometry

The relative abundance of total GDF15 was determined using the sum of H and D peptide. In studies of homozygote fetuses and heterozygous mothers (at position H202D), the proportion of fetal derived peptide in the maternal circulation was determined by calculating the proportion of discordant maternal peptide (discordant maternal peptide/total GDF15) and multiplying this by 2.

For pregnancies where the fetus was heterozygous, and the mother was homozygous for the reference allele (HH) at position H202D the proportion of the discordant fetal peptide was calculated by dividing this by total GDF15 and multiplying this by 2. Noticing that this produced nonsensical fetal proportions of GDF15 (e.g. in excess of 100%) in almost all samples tested, we calculated the average proportion of D-peptide in each pregnancy and used a one sample t-test to determine if the D-peptide constituted greater than 50% of total GDF15.

Linear mixed models with random intercepts implemented in the *LmerTest* package (https://cran.r-project.org/web/packages/lmerTest/index.html) were used to characterise the effect of gestational age on relative abundance of natural log transformed total circulating GDF15 measured by mass spectrometry.

### Functional studies of C211G

#### Plasmid construction

The expression vector for C-terminally Flag-tagged full-length human GDF15 was obtained from Genscript. The C211G mutant was generated by site-directed mutagenesis of the wild-type vector using the QuikChange II protocol (Agilent). To generate the Myc-tagged versions, the sequences corresponding to Flag tags were replaced by those encoding for Myc tags using the In-Fusion PCR cloning system (Takara) according to the kit’s guidelines. All plasmid sequences were confirmed by direct nucleotide sequencing.

#### Cell culture and transfection

Human embryonic kidney (HEK) 293T cells were obtained from ECACC/PHE and maintained in DMEM (Gibco) supplemented with 10% fetal bovine serum (Hyclone), 1% penicillin-streptomycin and 2 mM L-glutamine (Invitrogen), in an incubator at 37 °C in 5% CO_2_/95% O_2_ atmosphere. All cell lines were routinely tested as negative for mycoplasma contaminations using VenorGem Classic Mycoplasma Testing PCR Kit (Minerva Biolabs).

Cells were transiently transfected using Lipofectamine 3000 (Invitrogen) in 12-well plates with a total of 1000 ng DNA, as directed by the manufacturer.

#### Immunoblotting and co-immunoprecipitation

72-hour post-transfection, conditioned media samples were harvested, centrifuged, and proteins denatured under reducing conditions at 70 °C. For immunoblotting of intracellular proteins, cells were washed twice with cold PBS and lysed in M-Per Mammalian Protein Extraction Reagent (Thermo Scientific) supplemented with protease inhibitors. Whole cell extracts were sonicated and cleared by centrifugation and protein concentration estimated using the Bio-Rad DC protein assay kit (Bio-Rad Laboratories).

For co-immunoprecipitation experiments, Flag-tagged proteins were immunoprecipitated with anti-Flag magnetic agarose (Pierce Anti-DYKDDDDK Magnetic Agarose, ThermoFisher), according to the manufacturer’s protocol. Elution of bound proteins was performed with reducing SDS-PAGE sample buffer. Proteins were resolved by SDS-PAGE in NuPAGE Novex 4-12% Bis-Tris gels and transferred onto nitrocellulose membranes using the iBlot system (Invitrogen). Membranes were then blocked in 50 mM Tris-HCl, pH 7.6, 150 mM NaCl, 0.1% Tween-20 and 5% non-fat milk for 1 hour at room temperature and probed for 18 hours at 4 °C with antibodies specific for Flag tag (M2, SIGMA), Myc tag (9E10, sc-40, Santa Cruz Biotechnology), or calnexin (Cell Signaling Technology).

### Mouse Studies

In Cambridge, all mouse studies were performed in accordance with UK Home Office Legislation regulated under the Animals (Scientific Procedures) Act 1986 Amendment Regulations 2012 following ethical review by the University of Cambridge Animal Welfare and Ethical Review Body (AWERB).

Adult wild-type C57BL/6J male or female mice were purchased from Charles River (Charles River Ltd, Manston Rd, Margate, Kent, CT9 4LT) and kept under controlled light (12 h light:dark cycle (6:00 h:18:00 h), temperature (22 ± 1 °C) and humidity conditions with *ad libitum* access to food (RM3(E) Expanded Chow (Special Diets Services)) and water.

On the day of the experiment mice were divided into two weight and sex matched groups, single-housed and injected subcutaneously (s.c.) with either vehicle control (PBS) or GDF15 long-acting protein (FC-GDF15) provided by Pfizer Inc. under a material transfer agreement [15] at the dose of 0.01mg/kg (N=17, 12 male, 5 female, in Control and 19 in FC_GDF15 group, 13 male, 6 female). Food intake and body weight were measured daily. On the 4^th^ day, human recombinant GDF15 (hrGDF15, Cat# Qk017, Qkine) was administered via s.c. injection as a single dose in the afternoon (5pm). In all mice food intake and body weight were measured 16 hours after injection of hrGDF15. One cohort of mice (n=7-8 males per group) were sacrificed at 9 am the morning after the hrGDF15 injection, while the remainder went on to have food intake and body weight measured at 5pm.

Human GDF15 was measured using the human GDF15 ELISA (Cat#DY957, R&D Systems, BioTechne). In the mouse study - one female animal assigned to the control group (vehicle) was excluded due to failed subcutaneous injection with human recombinant GDF15. In addition, a food intake data point of another female vehicle control mouse (overnight food intake the day before treatment with human recombinant GDF15) was excluded from the analysis due to a transcription error during data collection. Hypothesis testing was conducted using repeated measures Two-way ANOVA or mixed effect models, with post-hoc Sidak’s test with the Geisser-Greenhouse correction implemented in Prism (Graphpad).

## Statistical analyses

Statistical analyses, including software employed, are described in the relevant sections of the text above.

## Acknowledgements

**HG Study:** We are grateful to Dr Rebecca McKay (NWA), the research midwives and staff of the early pregnancy admissions unit for their support to recruitment for the HG study. We are also grateful to the research midwives and the Daphne ward in Cambridge University Hospitals NHS Foundation Trust for recruitment of patients. We thank the women who participated in the HG study and the National Institute of Health Research (NIHR) Clinical Research Network (CRN Eastern) for supporting research midwives and nurses at study sites during this research study. This research was also supported by the National Institute for Health Research (NIHR) Cambridge Biomedical Research Centre (BRC) and the Core Biochemical Assay Laboratory (CBAL). The views expressed are those of the author(s) and not necessarily those of the NIHR or the Department of Health and Social Care. **Generation Scotland:** We are grateful to all the families who took part, the general practitioners and the Scottish School of Primary Care for their help in recruiting them, and the whole Generation Scotland team, which includes interviewers, computer and laboratory technicians, clerical workers, research scientists, volunteers, managers, receptionists, healthcare assistants and nurses. **CROATIA-Korcula:** We would like to acknowledge the contributions of the recruitment team in Korcula, the administrative teams in Croatia and Edinburgh and the people of Korcula. The exome sequencing was performed by the Regeneron Genetics Center. We thank the women who participated in the “Genes and risk factors for Hyperemesis Gravidarum” study and the families that participated in the “Fetal genes associated with recurrence of Hyperemesis Gravidarum” study. We would like to thank the research participants and employees of 23andMe for making this work possible. We would like to thank the Core Biochemical Assay Laboratory and the Proteomics and Peptidomics core at the Wellcome-MRC Institute of Metabolic Science for their services and technical expertise. We thank Roche and Ansh labs for the kind provision of reagents for GDF15 immunoassays. We thank Marko Hynoven for the kind gift of synthetic GDF15 peptides. We would like to thank Debra Rimmington for providing administrative support for the mouse studies.

For the purpose of open access, the author has applied a Creative Commons Attribution (CC BY) licence to any Author Accepted Manuscript version arising from this submission.

## Contribution statement

MF, NR, IC, SML, CP, GCSS, DSCJ, APC, CLM, SM, CH, NM and SOR designed the study. CP, IH, and KKO completed the work within the Cambridge Baby Growth Study. CP, ASC, MB and CLM undertook the HG study. AC, OP, GT and CH led the CROATIA-Korcula study. GDF15 was measured in the Cambridge Baby Growth Study and CROATIA-Korcula by PB and KB, and by CP, EC, GCSS and DSCJ in the HG study. Analysis of these studies was undertaken by CP and SML. Mass spectrometry studies were conducted by RGK and ALG and analysed by RGK, SML and DSCJ. SG genotyped participants in POPS at the H202D site using placental RNA-Seq data, confirmatory genotyping in umbilical cord DNA was done by DW and KR. NR conducted the *in vitro* experiments characterising the effects of the C211G variant. AC, PW, NS and CH conducted GDF15 pQTL discovery in Generation Scotland. SML, YBHL and NM conducted the common variant association analyses of GDF15 risk and HG including Mendelian randomization and colocalization analyses. NY, AP and SM conducted the Thalassemia studies. MF, VC, PM, KMG, EJ and AK conducted the studies of C211G fetal and maternal genotye on NVP. IC and APC conducted the mouse studies, IC and SML analysed the data, APC supervised the mouse studies. MF, NR, IC, SML, CP, RGK, GCSS, DSCJ, APC, CLM, SM, SH, NM and SOR wrote the manuscript, and all authors reviewed the manuscript for important intellectual content. This publication is the work of the authors, and MF, GCSS, DSCJ, APC, CLM, SM, SH, NM and SOR will serve as guarantors for the contents of this paper.

## Data availability

Summary statistics of the GDF15 GWAS in Generation Scotland can be obtained from the authors upon reasonable request. For the hyperemesis gravidarum GWAS: qualified researchers can contact to gain access to full GWAS summary statistics following an agreement with 23andMe that protects 23andMe participant privacy, directly relevant replication data are available upon request.

## Code availability

Code for data analysis is freely available upon request

