## Supplemental Tables for "Fetally-encoded GDF15 and maternal GDF15 sensitivity are major determinants of nausea and vomiting in human pregnancy"

**Supplementary Table 1** Recovery of synthetic GDF15 peptides by the MSD R&D DuoSet ELISA and the Ansh Total GDF15 ELISA.

| **Peptide** | **Theoretical** **pg/ml** | **MSD R&D DuoSet GDF15** | | | **Ansh Total GDF15** | | |
| --- | --- | --- | --- | --- | --- | --- | --- |
|  |  | **Analysed in 2021** | | | **Analysed in September 2022** | | |
|  |  | **Measured** | **Recovery to theoretical** | **Recovery  to HH** | **Measured** | **Recovery to theoretical** | **Recovery to HH** |
|  |  | **pg/ml** | **%** | **%** | **pg/ml** | **%** | **%** |
| HH | 2000 | 3329 | 166 | NA | 2270 | 114 | NA |
| HH | 667 | 1027 | 154 | NA | 677 | 101 | NA |
| HH | 222 | 367 | 165 | NA | 216 | 97 | NA |
| HH | 74 | 121 | 164 | NA | 81 | 109 | NA |
| HD | 2000 | 2384 | 119 | 72 | 2768 | 138 | 122 |
| HD | 667 | 795 | 119 | 77 | 829 | 124 | 124 |
| HD | 222 | 282 | 127 | 77 | 266 | 120 | 123 |
| HD | 74 | 88 | 119 | 73 | 90 | 121 | 111 |
| DD | 2000 | 1269 | 63 | 38 | 2009 | 100 | 89 |
| DD | 667 | 446 | 67 | 43 | 573 | 86 | 85 |
| DD | 222 | 155 | 70 | 42 | 164 | 74 | 76 |
| DD | 74 | 55 | 74 | 45 | 57 | 76 | 70 |

HH: Homodimeric peptide containing the reference H amino acid at position 6 of mature GDF15 (position 202 of the unprocessed form). HD: This is an admixture of synthetic peptides in the approximate ratio of HH:HD:DD of 1:2:1, akin to what would be expected in the circulation of heterozygote carriers of the H202D variant if all possible conformations of the peptide are secreted and eliminated equally. DD: Homodimeric peptide containing the alternate D amino acid at position 6 of mature GDF15. Recovery is expressed to the nearest %. The performance of the R&D DuoSet® assay adapted for use on the Meso Scale Discovery® (MSD) platform is presented for comparison [1].

**Supplementary Table 2** Characteristics of participants in the Cambridge Baby Growth Study

| **Characteristic** | **Reported No Nausea or Vomiting in Pregnancy (n=148)** | **Reported Vomiting in Pregnancy (n=168)** | **P-value** |
| --- | --- | --- | --- |
| Age (years) | 34.0 (33.2-34.6) | 32.9 (32.2-33.5) | 0.04 |
| Parity (n 0/1/2/>2) | 82/52/12/2 | 75/73/14/6 | 0.3 |
| Gestational age when GDF15 measured (weeks) | 14.9 (14.6-15.1) | 15.0 (14.8-15.2) | 0.4 |
| Pre-pregnancy BMI (kg/m^2^) | 23.8 (23.1-24.6) | 24.1 (23.5-24.8) | 0.5 |
| Reported taking anti-emetics in pregnancy   (n yes/no) | 0/148 | 18/150 | <0.001 |
| Reported smoking in pregnancy (n yes/no) | 5/143 | 9/159 | 0.4 |

Data are mean (95% confidence interval) or numbers. P-values are derived from linear regression models for continuous variables. Rates of anti-emetic usage and smoking status were compared between groups using Fisher’s exact and Chi-squared test, respectively.

**Supplementary Table 3** Serum GDF15 concentrations in women in the Cambridge Baby Growth study

| **Statistical Model** | **Serum GDF15 Concentrations (pg/mL)** | | **P-value** |
| --- | --- | --- | --- |
|  | **Women with no nausea or vomiting (n=148)** | **Women with vomiting  (n=168)** |  |
| Unadjusted | 13,198 (12,421-14,022) | 15,442 (14,588- 16,346) | 2.4x10^-4^ |
| Adjusted for gestational age | 13,181 (12,395-14,018) | 15,431 (14,573-16,339) | 2.5x10^-4^ |
| Adjusted for gestational age and BMI | 12,961 (12,161-13,813) | 15,542 (14,663-16,474) | 4.3x10^-5^ |

Data are geometric means (95% confidence intervals). P-values and adjusted GDF15 concentrations are derived from linear regression models of natural-log transformed GDF15.

**Supplementary Table 4**
Characteristics of cases and controls from the Hyperemesis Gravidarum study

| **Characteristic** | **Reported Minimal Nausea or Vomiting in Pregnancy (n=56)** | **Hyperemesis Gravidarum (n=57)** | **P-value** |
| --- | --- | --- | --- |
| Age (years) | 31.6 (30.1-33.1) | 29.7 (28.2-31.2) | 0.09 |
| Pregnancy length when  GDF15 measured (weeks) | 10.1 (8.9-11.4) | 9.0 (7.8-10.2) | 0.2 |
| Pre-pregnancy BMI (kg/m^2^) | 24.8 (22.2-27.3) | 25.1 (23.7-26.6) | 0.8 |
| Reported taking anti-emetics in current pregnancy  (n yes/no) | 0/56 | 38/19 | <0.001 |
| Reported requiring rehydration in current pregnancy (n yes/no) | 1/55 | 23/34 | <0.001 |
| “Current Nausea” Score at time of sampling (out of 10) | 0.2 (0-0.7) | 6.0 (5.5-6.5) | 3.6x10^-31^ |
| “Worst Nausea” in pregnancy Score (out of 10) | 1.6 (1.2-2.0) | 9.4 (9.0-9.8) | 5.0x10^-51^ |
| “Current Vomiting” Score at time of sampling (out of 10) | 0 (0-0.7) | 3.5 (2.9-4.2) | 2.1x10^-11^ |
| “Worst Vomiting” in pregnancy Score (out of 10) | 0.1 (0-0.5) | 8.7 (8.3-9.1) | 1.0x10^-54^ |
| Sum of Nausea and Vomiting Scores (out of 40) | 1.9 (0.7-3.0) | 27.6 (26.5-28.7) | 1.1x10^-57^ |
| Reported smoking in pregnancy (n yes/no) | 3/53 | 11/46 | 0.04 |

Data are mean (95% confidence interval) or numbers. P-values are from linear regression for continuous variables and Fisher’s exact test for categorical variables.

**Supplementary Table 5**
Serum GDF15 concentrations in women in the hyperemesis gravidarum study

| **Statistical Model** | **Serum GDF15 Concentrations (pg/mL)** | | **P-value** |
| --- | --- | --- | --- |
|  | **Women with minimal   nausea or vomiting  (n=56)** | **Women with HG  (n=57)** |  |
| Unadjusted | 9,396 (7,981-11,062) | 13,172 (11,204-15485) | 4.3x10^-3^ |
| Adjusted for gestational age | 8,899 (7,674-10,320) | 13,314 (11,512-15,399) | 2.1x10^-4^ |
| Adjusted for gestational age and BMI | 8,696 (5,461-13,848) | 10,079 (7,810-13,006) | 0.01 |
| Adjusted for gestational age and smoking status | 8,630 (7,448-9,999) | 13,595 (11,778-15,692) | 3.3x10^-5^ |

Data are geometric mean (95% confidence interval). P-values and adjusted GDF15 concentrations are derived from linear regression models of natural-log transformed GDF15.

**Supplementary Table 6**
Genotyping results obtained using placental RNAseq data to select samples used for MS-based measurement of GDF15 in maternal plasma during pregnancy. Fetal genotypes were confirmed by PCR using umbilical cord DNA as described in the methods.

| **Maternal age at enrolment** | **Ref.allele** | **Alt.allele** | **RefCount** | **AltCount** | **Fetal**  **Genotype** | **Maternal**  **Genotype** |
| --- | --- | --- | --- | --- | --- | --- |
| 32 | C | G | 42 | 49 | C/G | C/C |
| 22 | C | G | 55 | 63 | C/G | C/C |
| 30 | C | G | 71 | 46 | C/G | C/C |
| 24 | C | G | 65 | 56 | C/G | C/C |
| 31 | C | G | 63 | 57 | C/G | C/C |
| 32 | C | G | 64 | 51 | C/G | C/C |
| 26 | C | G | 65 | 58 | C/G | C/C |
| 24 | C | G | 36 | 46 | C/G | C/C |
| 28 | C | G | 41 | 44 | C/G | C/C |
| 27 | C | G | 41 | 64 | C/G | C/C |
| 29 | C | G | 47 | 41 | C/G | C/C |
| 37 | C | G | 68 | 58 | C/G | C/C |
| 39 | C | G | 0 | 118 | G/G | C/G |
| 31 | C | G | 1 | 101 | G/G | C/G |
| 29 | C | G | 0 | 100 | G/G | C/G |
| 36 | C | G | 2 | 85 | G/G | C/G |
| 25 | C | G | 2 | 29 | G/G | C/G |
| 20 | C | G | 1 | 26 | G/G | G/G |
| 32 | C | G | 129 | 0 | C/C | C/C |
| 34 | C | G | 126 | 0 | C/C | C/C |
| 30 | C | G | 125 | 0 | C/C | C/C |
| 29 | C | G | 125 | 0 | C/C | C/C |
| 28 | C | G | 124 | 0 | C/C | C/C |
| 25 | C | G | 121 | 0 | C/C | C/C |
| 31 | C | G | 116 | 0 | C/C | G/C |
| 25 | C | G | 110 | 0 | C/C | G/C |

**Supplementary Table 8**. The effect of fetal genotype on nausea and vomiting in pregnancy in mothers carrying the C211G variant in *GDF15*.

| **Mother**  **TG (C211G)** | **Child** | **Child**  **Genotype** | **Mother**  **HG** | **Prescription  Antiemetic** | **IV**  **Fluids** | **Emergency**  **Room** | **Hospitalized** | **Symptom**  **Resolution** |
| --- | --- | --- | --- | --- | --- | --- | --- | --- |
| 1 | 1.1 | TT | Y | Y | Y | Y | Y | at birth |
|  | 1.2 | TT | Y | Y | Y | Y | Y | at birth |
| 2 | 2.1 | TG | N | N | N | N | N | T1 |
|  | 2.2 | TT | Y | Y | Y | Y | Y | T3 |
| 3 | 3.1 | TG | Y | Y | Y | Y | N | T2 |
|  | 3.2 | TT | Y | Y | Y | Y | N | T3 |
|  | 3.3 | TG | Y | Y | Y | Y | N | after birth |
|  | 3.4 | TT | Y | Y | N | N | N | after birth |
|  | 3.5 | TT | Y | Y | Y | Y | N | after birth |
| 4 | 4.1 | TG | Y | Y | Y | N | N | after birth |
|  | 4.2 | TG | Y | Y | Y | N | N | after birth |
|  | 4.3 | TT | Y | Y | Y | N | Y | after birth |
|  | 4.4 | TT | Y | Y | N | N | N | after birth |
| 5 | 5.1 | TT | Y | Y | Y | Y | N | T2 |
|  | 5.2 | TT | Y | Y | Y | Y | Y | T2 |
|  | 5.3 | TG | N | N | N | N | N | T1 |
| 6 | 6.1 | TG | N | N | N | N | N | T1 |

C211G (TG) Mothers have Hyperemesis Gravidarum with all 10 pregnancies carrying a homozygous TT fetus, but normal NVP (no treatment) in 3/7 pregnancies carrying a TG fetus.  (T1,2,3=Trimester 1, 2, or 3). IV fluids – Intravenous Fluids.

**Supplementary Table 9** Mendelian Randomization of circulating GDF15 levels on hyperemesis gravidarum risk using the Roche assay

| **Method** | **Slope/Intercept** | **Estimate** | **Std Error** | **95% CI** | | **P-value** |
| --- | --- | --- | --- | --- | --- | --- |
| Simple median | slope | -0.79 | 0.05 | -0.88 | -0.70 | 8.47E-66 |
| Weighted median | slope | -0.95 | 0.04 | -1.03 | -0.87 | 1.55E-113 |
| Penalized weighted median | slope | -0.95 | 0.04 | -1.03 | -0.87 | 1.56E-113 |
| IVW | slope | -0.35 | 0.04 | -0.43 | -0.27 | 6.98E-17 |
| Penalized IVW | slope | -0.35 | 0.04 | -0.43 | -0.27 | 6.98E-17 |
| Robust IVW | slope | -0.35 | 0.04 | -0.43 | -0.27 | 6.98E-17 |
| Penalized robust IVW | slope | -0.35 | 0.04 | -0.43 | -0.27 | 6.98E-17 |
| MR-Egger | slope | -0.33 | 0.04 | -0.41 | -0.25 | 2.00E-15 |
| MR-Egger | intercept | 0.00 | 0.00 | -0.01 | 0.00 | 2.94E-02 |
| Penalized MR-Egger | slope | -0.33 | 0.04 | -0.41 | -0.25 | 2.00E-15 |
| Penalized MR-Egger | intercept | 0.00 | 0.00 | -0.01 | 0.00 | 2.94E-02 |
| Robust MR-Egger | slope | -0.33 | 0.04 | -0.41 | -0.25 | 2.00E-15 |
| Robust MR-Egger | intercept | 0.00 | 0.00 | -0.01 | 0.00 | 2.94E-02 |
| Penalized robust MR-Egger | slope | -0.33 | 0.04 | -0.41 | -0.25 | 2.00E-15 |
| Penalized robust MR-Egger | intercept | 0.00 | 0.00 | -0.01 | 0.00 | 2.94E-02 |

We estimated putative causal effects (ie slope) of circulating GDF15 levels in the non-pregnant state on hyperemesis gravidarum (HG) risk using m=259 variants from Roche-based pQTL summary data (n=18,184) and 23andMe HG summary data (n=17,062). Harmonized pQTL, GWAS, and LD estimated from UK Biobank WGS individuals. The causal effect estimates represent the change in HG risk in log-odds per standard deviation increase in circulating GDF15.

**Supplementary Table 10** Mendelian Randomization of circulating GDF15 levels on hyperemesis gravidarum risk are robust to LD reference

| **Method** | **LD Reference** | **Estimate** | **Std Error** | **95% CI** | | **P-value** |
| --- | --- | --- | --- | --- | --- | --- |
| IVW | UKBB | -0.35 | 0.04 | -0.43 | -0.27 | 6.98E-17 |
| IVW | 1000G | -0.36 | 0.03 | -0.42 | -0.30 | 9.41E-30 |

To assess the stability of our results to choice of LD reference panel, we re-estimated putative causal effects (ie slope) of circulating GDF15 levels in the non-pregnant state on hyperemesis gravidarum (HG) risk using Roche-based pQTL summary data (n=18,184) and 23andMe HG summary data (n=17,062). However, LD was estimated from 1000G WGS individuals which resulted in m=310 variants (see Methods). The causal effect estimates represent the change in HG risk in log-odds per standard deviation increase in circulating GDF15.

**Supplemental Table 11** Mendelian Randomization of circulating GDF15 levels on hyperemesis gravidarum risk are robust to the protein altering variant p.H202D (rs1058587)

| **Method** | **Analysis** | **Estimate** | **Std Error** | **95% CI** | | **P-value** |
| --- | --- | --- | --- | --- | --- | --- |
| IVW | Marginal | -0.35 | 0.04 | -0.43 | -0.27 | 6.98E-17 |
| IVW | Conditional | -0.21 | 0.02 | -0.25 | -0.16 | 3.82E-19 |

To assess the stability of our results to variant rs1058587, which was previously suggested to confound quantification [2, 3], we re-estimated putative causal effects (ie slope) of circulating GDF15 levels in the non-pregnant state on hyperemesis gravidarum (HG) risk using m=258 variants from Roche-based pQTL summary data (n=18,184) and 23andMe HG summary data (n=17,062) after conditioning (ie residualizing) on variant rs1058587 (see Methods). Results from IVW on reported pQTL/GWAS data are listed under "Marginal", while updated MR results obtained from a conditional analysis are listed under "Conditional". The causal effect estimates represent the change in HG risk in log-odds per standard deviation increase in circulating GDF15.

**Supplementary Table 12** Colocalization analysis identifies two shared genetic signals for circulating GDF15 and hyperemesis gravidarum risk

| **hit.pQTL** | **hit.GWAS** | **PP.H0.abf** | **PP.H1.abf** | **PP.H2.abf** | **PP.H3.abf** | **PP.H4.abf** |
| --- | --- | --- | --- | --- | --- | --- |
| rs16982345 | rs45543339 | 7.14E-231 | 1.72E-14 | 2.75E-219 | 4.63E-03 | 9.95E-01 |
| rs1227734 | rs1227731 | 1.07E-189 | 3.75E-04 | 2.09E-188 | 5.31E-03 | 9.94E-01 |

Multi-SNP colocalization analyses was performed using the R package coloc (see Methods). PP.HX.abf corresponds to the posterior probability that a variant cluster is null (X=0), is private to GDF15 levels (X=1), hyperemesis gravidarum (HG) risk (X=2), contributes independently to GDF15 levels and HG risk (X=3), and colocalizes across both traits (X=4), with a posterior probability of >0.8 generally considered as strong evidence of colocalization.

**Supplementary Table 13** 
The prevalence of nausea, vomiting and loss of appetite of females with thalassaemia

| **Parameter** | **Thalassaemia group (n=20)** | **Non-thalassaemia group (n=20)** | **Adjusted odds ratio (95% CI)** | **P-value** |
| --- | --- | --- | --- | --- |
| Nausea | 1 (5%) | 12 (60%) | 0.026 (0.002-0.310) | 0.004^#^ |
| Vomiting | 1 (5%) | 13 (65%) | 0.021 (0.002-0.248) | 0.002^#^ |
| Loss of appetite | 3 (15%) | 13 (65%) | 0.066 (0.010-0.443) | 0.005^#^ |
| Nausea and vomiting persistent beyond the first trimester | 1 (5%) | 3 (15%) | 0.345 (0.024-4.987) | 0.435^#^ |
| Nausea and vomiting persistent throughout pregnancy | 0 | 1 (5%) | - | 1.00* |
| Nausea and vomiting requiring treatment | 0 | 6 (30%) | - | 0.020* |
| Nausea and vomiting requiring hospitalisation | 0 | 3 (15%) | - | 0.231* |

Non-thalassaemia group: control group without thalassemia, matched for age and ethnicity. The prevalence of symptoms is adjusted for parity, number of children and time since index pregnancy. ^#^ Logistic regression  * Fisher’s exact test
