## Supplemental Figures for "Fetally-encoded GDF15 and maternal GDF15 sensitivity are major determinants of nausea and vomiting in human pregnancy"

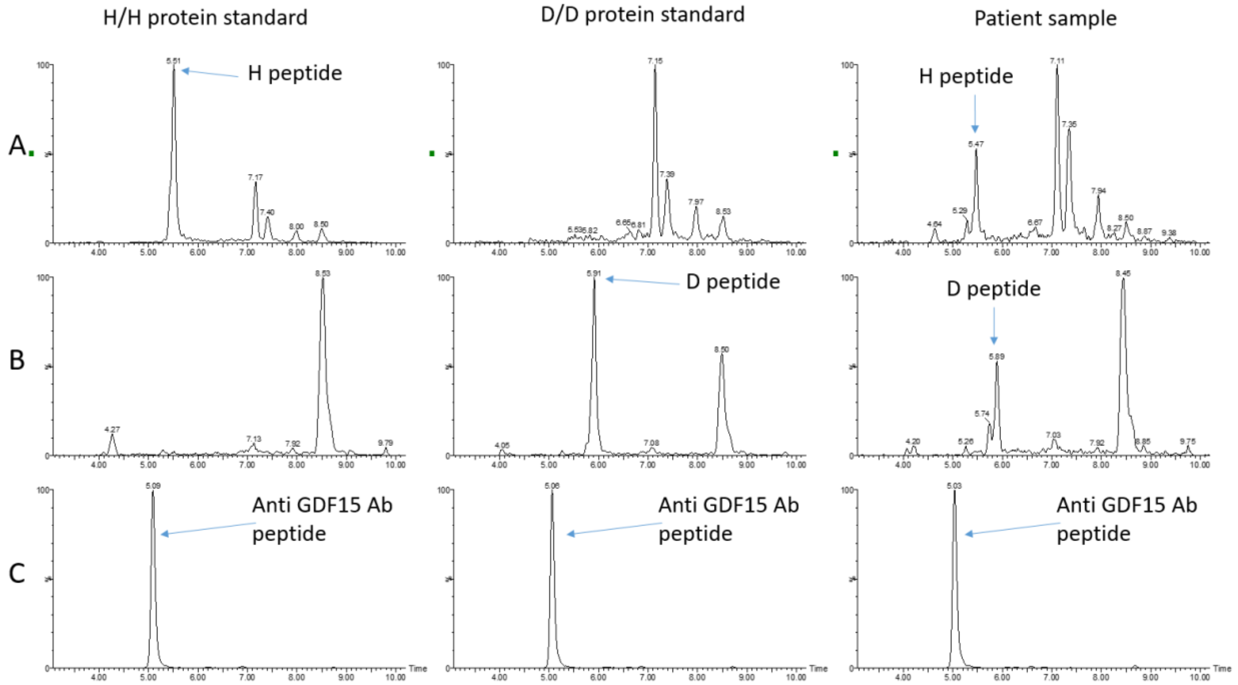


**Extended Data Figure 1. LC-MS/MS traces of two GDF15 related peptides and the murine anti-GDF15 antibody peptide from the heterozygous fetus analysis. A:** N-terminal peptide from the wild-type protein, RT=~5.47. **B**: N-terminal peptide from the mutant protein, RT=~5.51. **C**: Peptide from the murine anti GDF15 antibody, RT=~5.09. Data shown is traces generated from extracted plasma spiked with mutant homodimer, wild type homodimer and an extracted patient sample.

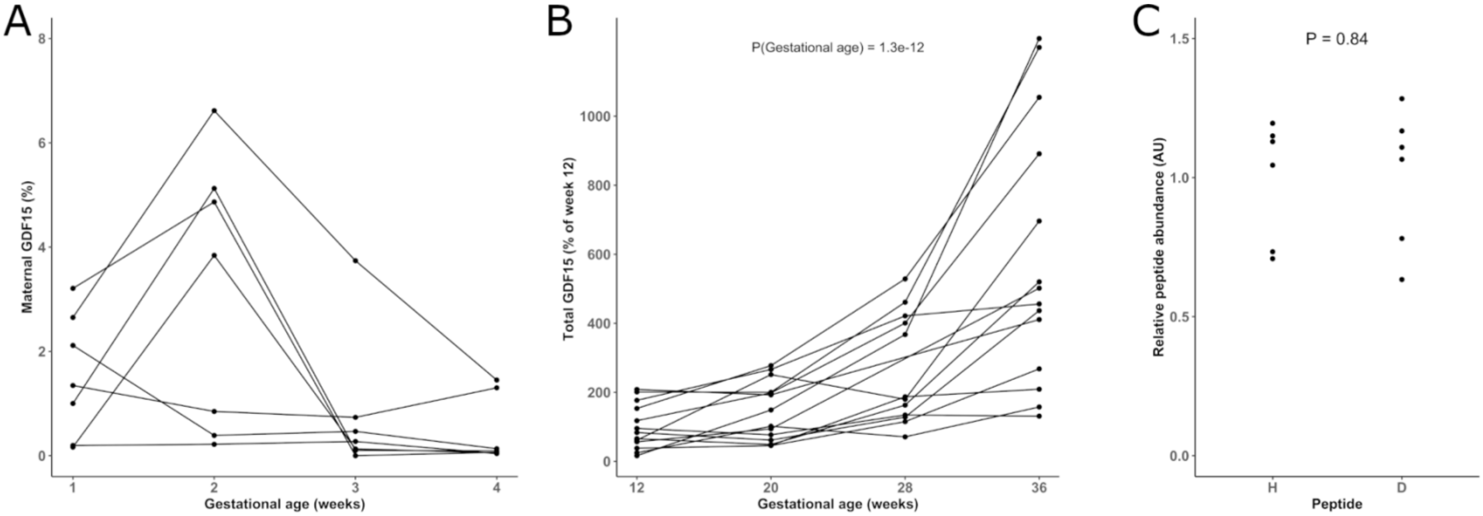


**Extended Data Figure 2. Measurement of fetal and maternally derived GDF15 in pregnancy.**
**A:** The estimated proportion of maternally derived GDF15 in 7 different pregnancies across 4 gestational ages where the fetus is homozygous for either the H or D at H202D and the mother is heterozygous at this site**. B:** The relative abundance of Total GDF15 measured by mass spectrometry in 14 different pregnancies where the fetus is homozygous for either H or D at H202D across 4 gestational ages, including the 7 genotype-discordant pregnancies presented in panel (A) and a further 7 pregnancies where the maternal genotype is concordant with the fetal genotype. Total GDF15 is expressed as a percentage of the mean value at 12 weeks gestation. P-value derived from a linear mixed model of log transformed GDF15 with random intercepts. **C:** The relative abundance of N-terminal peptides from synthetic GDF15 homodimers with H or D at position 202 extracted using the R&D anti-GDF15 capture antibody coupled to magnetic beads. Plasma was fortified at the same concentration for each protein, extracted (n=6) and analyzed by Orbitrap MS. AU = arbitrary units.

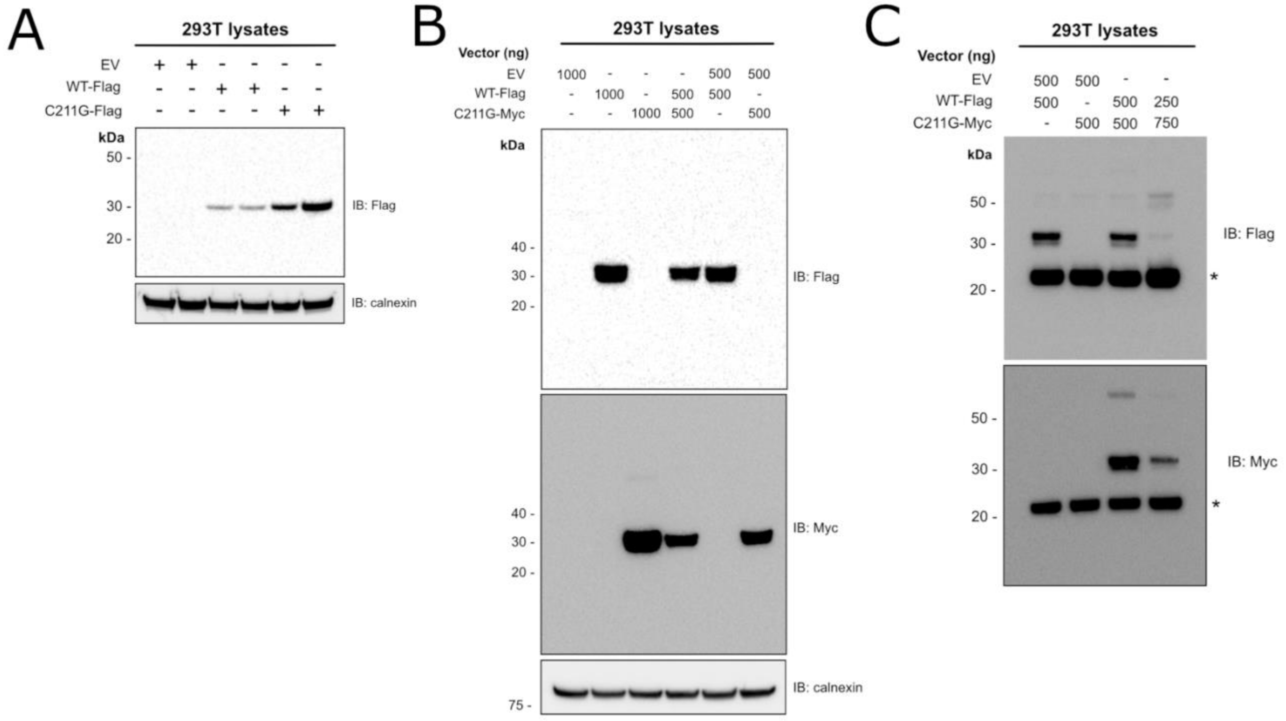


**Extended Data Figure 3. The C211G mutant is expressed intracellularly and heterodimerizes with its wild-type counterpart. A:** Western blotting of cell lysates expressing Flag-tagged fusions of wild-type GDF15 (WT-Flag) or GDF15 C211G (C211G-Flag). **B:** Co-expression of wild-type GDF15 (WT-Flag) and Myc-tagged GDF15 C211G (C211G-Myc) does not impair the intracellular expression of wild-type GDF-15. **C:** WT and C211G form intracellular heterodimers, as judged by the co-immunoprecipitation of WT-Flag and C211G-Myc using anti-Flag antibodies. The asterisk marks co-eluted Immunoglobulin light chains. Replicates, N=3, representative images are shown. EV indicates transfections with the empty plasmid backbone only.

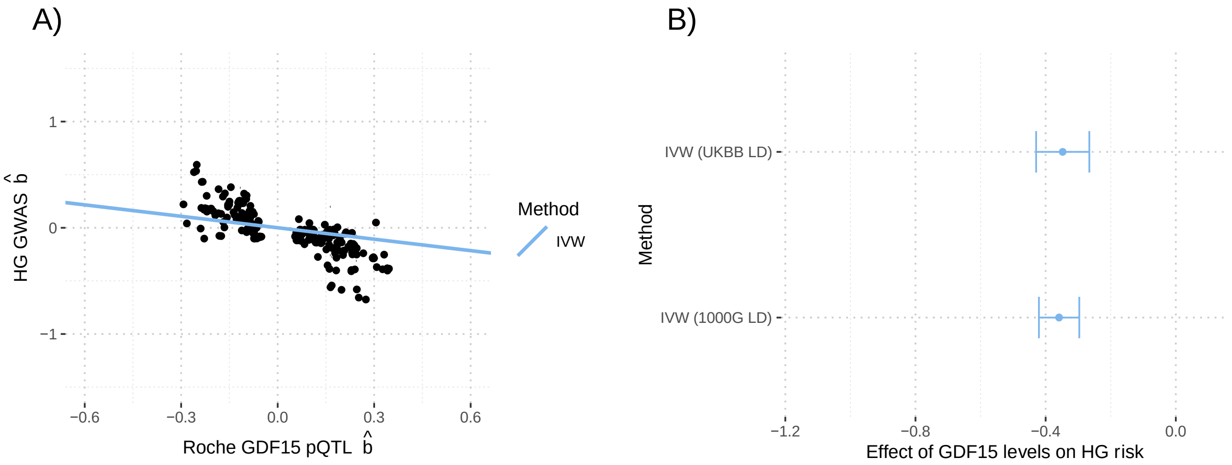


**Extended Data Figure 4. Mendelian Randomization estimates are robust to LD reference panel.** MR was performed using m=259 SNPs with genome-wide evidence of pQTL effects on GDF15 levels within 1Mb GDF15 locus and adjusted using LD estimates from 1000G WGS individuals (n=489; see Methods). **A:** Scatterplot of HG GWAS effect estimates (ie log-odds) vs Roche-based GDF15 pQTL effect estimates. Vertical and horizontal lines represent 95% confidence intervals of HG effects and GDF15 effects, respectively. Causal effects were estimated using LD-aware IVW MR and depicted as a regression line. **B:** Forest plot of the IVW MR causal effect-size estimates of circulating GDF15 levels on HG risk from UK Biobank and 1000G LD references. Each point represents the estimated causal effect and 95% confidence interval of a 1 standard deviation increase in circulating GDF15 in the non-pregnant state on HG risk in log-odds. The null of no mediating/causal effect is represented as a solid red line at 0.

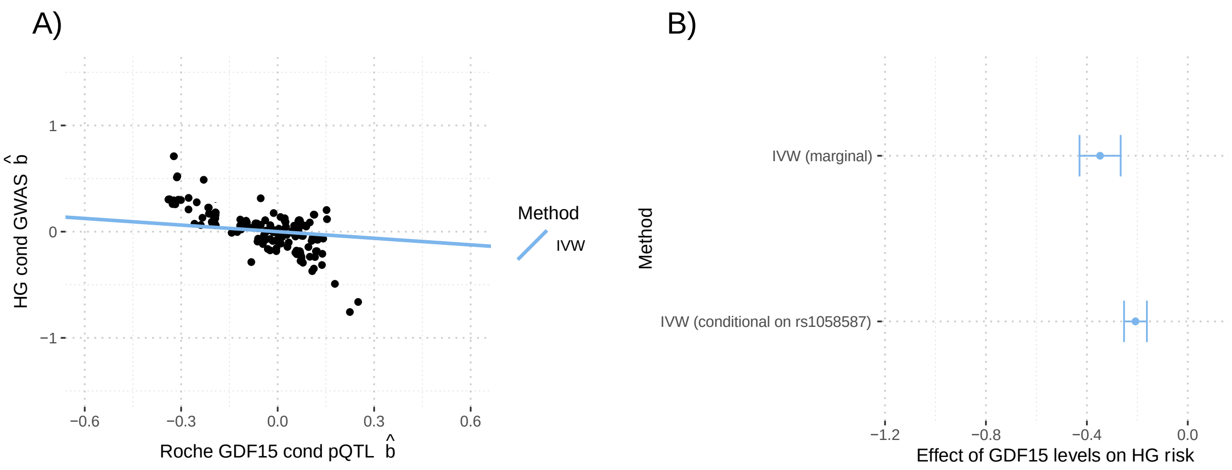


**Extended Data Figure 5. Mendelian Randomization estimates are robust to previously reported confounder SNP.** MR was performed using m=258 SNPs with genome-wide evidence of pQTL effects on GDF15 levels within 1Mb GDF15 locus after residualizing (ie conditioning) on the effect of variant rs1058587, which was previously suggested to confound quantification of GDF15 levels [1, 2]. Results were adjusted using LD estimates from UKBiobank WGS individuals (n=138335; see Methods). **A** Scatterplot of conditional HG GWAS effect estimates (ie log-odds) vs conditional Roche-based GDF15 pQTL effect estimates. Vertical and horizontal lines represent 95% confidence intervals of HG effects and GDF15 effects, respectively. Causal effect estimates obtained using LD-aware IVW MR and reflected as regression lines. **B:** Forest plot of the causal effect-size estimates of circulating GDF15 levels on HG risk from standard (ie marginal) pQTL/GWAS results and those obtained using pQTL/GWAS results conditioned on variant rs1058587. Each point represents the estimated causal effect and 95% confidence interval of a 1 standard deviation increase in circulating GDF15 in the non-pregnant state on HG risk in log-odds. The null of no mediating/causal effect is represented as a solid red line at 0.

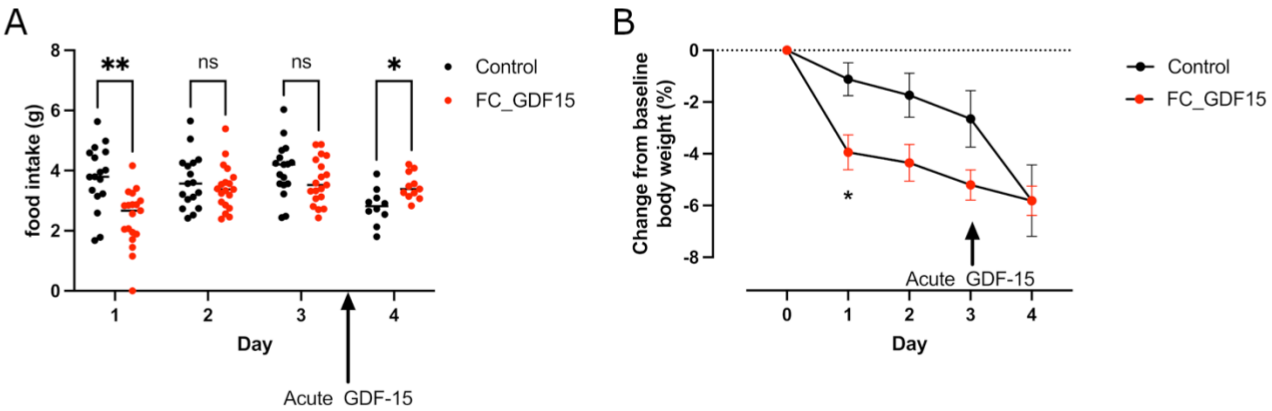


**Extended Data Figure 6. Longitudinal effects of long acting GDF15 on food intake and body weight and mice**

**A-B:** The effects of 0.01mg/kg of Fc-GDF15-15 fusion protein (FC_GDF15) or vehicle control (PBS) on food intake **(A)** or body weight **(B)**. In (A) Days 1 – 3 represent 24-hour food intake from 5pm to 5pm after treatment with control or FC_GDF15. Day 4 represents food intake from 5pm to 5pm after both groups received an acute bolus of human recombinant GDF15 (0.1mg/kg). In (B) change in body weight at 5pm is presented as a percentage of baseline body weight. Days 1-3: N=17 (12 male, 5 female) in Control and 19 in FC_GDF15 group (13 male, 6 female). Day 4: N=10 (5 male, 5 female) in Control and 11 in FC_GDF15 group (5 male, 6 female) – as one cohort of mice were sacrificed at 9am on Day 4. Hypothesis testing was conducted using a mixed-effects model. Post-hoc testing comparing Control and FC_GDF15 treated groups was undertaken with the Sidak test to correct for multiple testing. *P<0.05, **P<0.01, ns = non-significant, P>0.05.
